# APMAT analysis reveals the association between CD8 T cell receptors, cognate antigen, and T cell phenotype and persistence

**DOI:** 10.1101/2025.01.08.631993

**Authors:** Jingyi Xie, Daniel G. Chen, William Chour, Rachel H. Ng, Rongyu Zhang, Dan Yuan, Jongchan Choi, Michaela McKasson, Pamela Troisch, Brett Smith, Lesley Jones, Andrew Webster, Yusuf Rasheed, Sarah Li, Rick Edmark, Sunga Hong, Kim M. Murray, Jennifer K. Logue, Nicholas M. Franko, Christopher G. Lausted, Brian Piening, Heather Algren, Julie Wallick, Andrew T. Magis, Kino Watanabe, Phil Mease, Philip D. Greenberg, Helen Chu, Jason D. Goldman, Yapeng Su, James R. Heath

## Abstract

Elucidating the relationships between a class I peptide antigen, a CD8 T cell receptor (TCR) specific to that antigen, and the T cell phenotype that emerges following antigen stimulation, remains a mostly unsolved problem, largely due to the lack of large data sets that can be mined to resolve such relationships. Here, we describe Antigen-TCR Pairing and Multiomic Analysis of T-cells (APMAT), an integrated experimental-computational framework designed for the high-throughput capture and analysis of CD8 T cells, with paired antigen, TCR sequence, and single-cell transcriptome. Starting with 951 putative antigens representing a comprehensive survey of the SARS-CoV-2 viral proteome, we utilize APMAT for the capture and single cell analysis of CD8 T cells from 62 HLA A*02:01 COVID-19 participants. We leverage this unique, comprehensive dataset to integrate with peptide antigen properties, TCR CDR3 sequences, and T cell phenotypes to show that distinct physicochemical features of the antigen-TCR pairs strongly associate with both T cell phenotype and T cell persistence. This analysis suggests that CD8+ T cell phenotype following antigen stimulation is at least partially deterministic, rather than the result of stochastic biological properties.

## Introduction

CD8 T cells play a major role in adaptive immunity against pathogens, exhibiting a functional diversity that includes, as major phenotypes, naïve, memory, effector memory, effector, and exhausted, with each of those phenotypes encompassing multiple sub- phenotypes^1,2^. Recent literature has suggested the existence of relationships between a given antigen-specific T cell clonotype and the phenotypic trajectory that clonotype can take following antigen stimulation. For example, tissue-resident, antigen-specific T cell clonotypes in both tumor and chronic viral infection settings have been intimately associated with specific phenotype differentiation trajectories^3–5^. We and others have shown that, for at least certain immunogenic viral antigens, T cell clonotype is the dominant factor in determining T cell phenotype^6–8^. These similar results, from tissue and peripheral blood, and in very different disease settings, suggest that there may be rules that underlie T cell clonotype-T cell phenotype relationships. Elucidating such relationships requires experimental methods for collecting a large, longitudinal data set in which the transcriptome-level phenotypes of antigen-specific T cells are co-measured across a large number and diversity of antigens, coupled with computational methods for elucidating what, if any, relationships between the T cell receptor (TCR) α/β genes, cognate peptide antigen – major histocompatibility complex (pMHC), and T cell phenotype exist.

Initial hints at what such a rule set might contain can be found in the literature. For example, the importance of hydrophobicity at certain TCR residues is known to associate with cytotoxicity and self-reactivity in CD4 T cells^6,9–11^, and the hydrophobicity of certain residues is known to associate with the immunogenicity of antigens^12^. These observations suggest that the biochemical nature of the TCR:pMHC interface may play an important roles in determining T cell phenotype and cell fate trajectory. This question is not just of fundamental importance to T cell immunology, but it is also highly relevant to the engineering of T cells for use as therapeutic agents for treatments of both cancers^13^ and autoimmune diseases^14,15^. However, existing studies predominantly explore only one aspect of the Antigen-TCR-phenotype interplay: either resolving the antigen specificity of T cells from biological samples^16–18^, or conducting T cell receptor analysis on existing dataset lacking antigen specificity (or containing only limited number of antigens)^6,19,20^. Recent advances highlight the need of high-throughput, high-dimensional approaches as powerful tools for identifying and analyzing antigen-specific CD8 T cells^17,20–22^.

We describe an integrated experimental-computational framework, APMAT for the antigen-specific capture of T cells from many patient biospecimens in parallel, and is integrated with single-cell multi-omics profiling designed to integrate paired antigen and TCR sequence with T cell phenotype. For antigen-specific T cell capture, we utilize large libraries of single-chain-trimers (SCTs)^22,23^. We have recently reported on the feasibility of these libraries to capture and characterize both viral antigen-specific and (tumor-associated) neoantigen-specific T cell populations. Here we use SCTs to construct a 951-element pMHC library that represents a comprehensive survey of putative Class I antigens, presented by HLA A*02:01, from across the full SARS-CoV-2 genome. The DNA-barcoded SCT-pMHC library (for n = 558 expressed SCT) is used to identify and, through single cell (sc) RNA-seq analysis, characterize SARS-CoV-2 specific T cells from 62 HLA A*02:01 participants at the stages of acute and post-acute COVID-19. We identify several viral antigen-specific T cell populations observed in previous work by ourselves and others^24–27^ demonstrating the feasibility of this framework on a whole viral proteome scale. Moreover, resultant data set allows for an in-depth exploration to reveal insights into how the physicochemical properties of the TCR-pMHC interface associate with T cell phenotype and T cell persistence. This study elucidates, for a whole viral proteome and a single HLA allele, the physiochemical basis linking TCR:pMHC molecular interactions to the phenotypic behavior of antigen-specific CD8+ T cells, and thus advances our understanding of immunological mechanisms.

## Results

### APMAT enables integrated multi-modal analysis of antigen-specific CD8 T cells

We utilized APMAT to identify SARs-CoV-2 antigen-specific T cell responses to COVID-19 infection in a previously described longitudinal cohort of 209 COVID-19 participants^25,26^. From this cohort, we selected 62 HLA- A*02:01 participants representing a range of COVID-19 disease severities (Fig. 1a) with longitudinal reference datasets available at acute (diagnosis and approximately 1-week post-diagnosis), and post-acute (2-3 months post-diagnosis) timepoints (Supplementary Table 1)^25,26^.

**Fig. 1:**
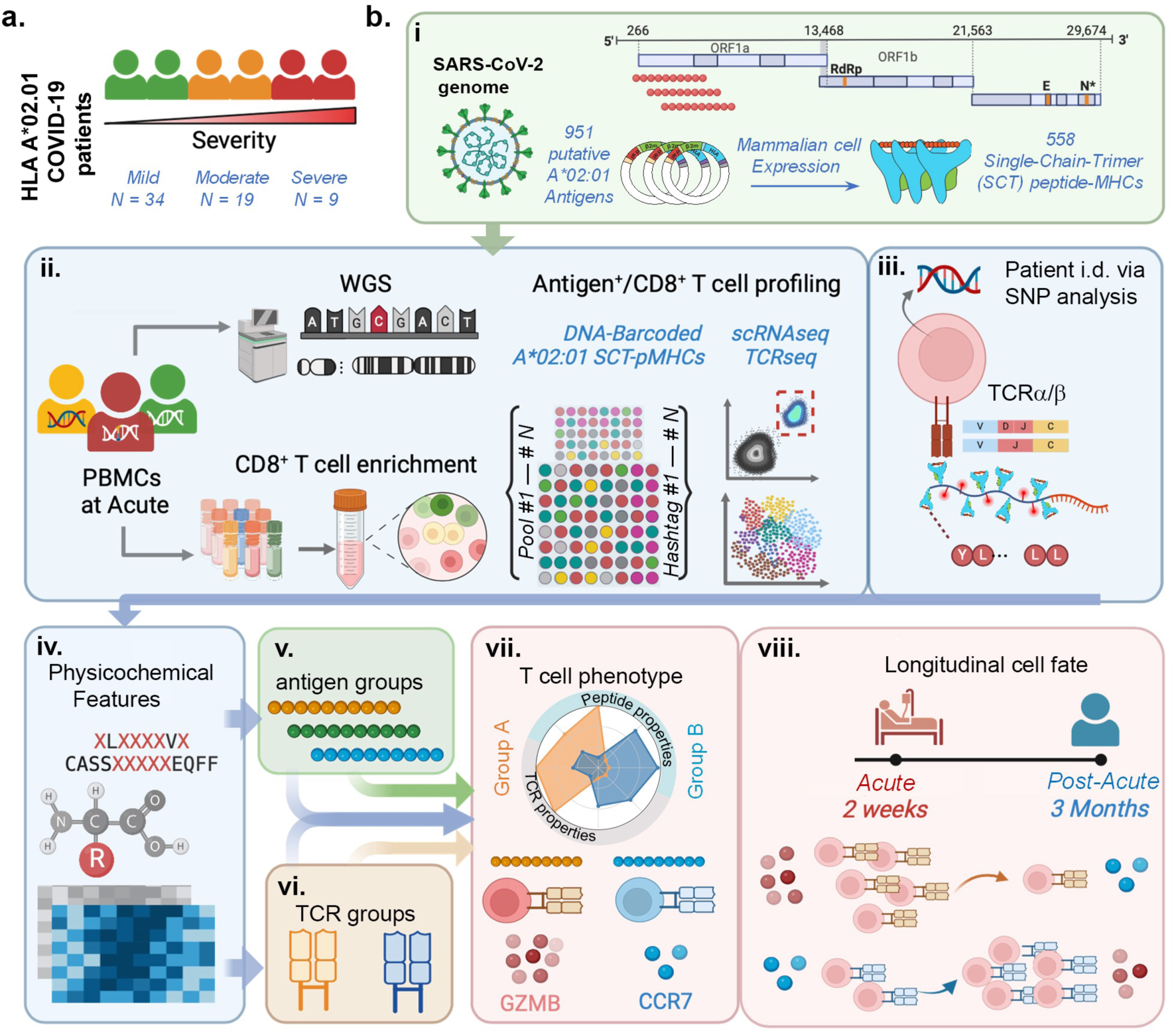
Schematic overview of antigen-TCR pairing and multiomic analysis of T-cells (APMAT) in COVID-19 participants a. Participants in the study, broken down by COVID-19 severity. b. Experimental and computational flow of APMAT. i. Construction of the large SCT-pMHC library representing HLA-A*02:01 peptide-MHC complexes covering antigens from across the full SARS-CoV- 2 proteome. ii. Antigen-specific T cells are captured from pooled patient PBMCs with barcoded SCT multimers for scRNAseq analysis. iii. Patient i.d.’s are assigned to each single cell by matching SNP analysis from whole genome with scRNAseq data. iv-vii. For each T cell, the physicochemical properties of the peptide antigen and the CDR3β domain of the TCRs are analyzed to identify statistical associations between the CDR3β-peptide antigen interface with T cell phenotype. viii. T cell clonotype persistence from acute disease to convalescence is similarly associated with the physicochemical properties of the TCR:antigen.

For antigen-specific CD8 T cell capture, we utilized DNA-barcoded, pMHC-like SCT multimers (SCT-dextramers). For library construction, we first utilized NetMHCpan^28^ to analyze the full SARS-CoV-2 viral genome (original strain) to resolve a list of 951 putative 9 - 10 mer antigens (Supplementary Table 2.1) for HLA-A*02:01. This peptide list was converted into PCR-optimized DNA primers and used to build the plasmid library. Plasmids were then transfected into Expi293 cells over four days to induce secretion of the SCT protein product. 558 of the 951 putative antigens yielded usable SCT product (Fig. 1b top). Each SCT was then purified, biotinylated, and assembled into a fluorophore-labeled and DNA-barcoded dextramer, using previously reported protocols^23^. To minimize batch effects and enable integrated analysis, PBMCs from all 62 participants, collected at the stage of acute disease, were pooled and profiled in parallel within one experiment (Fig. 1b middle left). SCT-dextramer-positive CD8 T cells were sorted for scRNAseq to profile gene expression, TCRα/β genes, and antigen specificity (Fig. 1b middle right). For each cell captured, antigen-specificity was identified through the dominant pool hashtag (Supplementary Fig. 1a) and dextramer (Supplementary Fig. 1b). Each cell was associated with a specific patient through comparison of de novo computed single nucleotide polymorphisms (SNPs) from scRNA-seq with germline whole genome sequence (WGS) profiles of all patients (Supplementary Fig. 1c). Sex gene validation further confirmed patient assignments (Supplementary Fig. 1d). Out of 19,103 cells from scRNA-seq data, only 166 (0.87%) were not assignable to a donor (Supplementary Table 2.2). With such a multi-modular dataset, the characteristics of the peptide antigens and the TCR CDR3 sequences can be integrated with T cell phenotype and longitudinal multi-omics datasets (Fig. 1b bottom).

### APMAT enables high-throughput representation of whole SARS-CoV-2 genome

We graph the distribution of putative antigens against the SARS-CoV-2 genome, and highlighted the 558 (58.7%) constructs that were expressed and used for the T cell capture experiment, and the 102 (18.3%) that captured at least 1 cell (Fig. 2a, Supplementary Fig. 2a). While APMAT identified CD8 T cells against antigens from across the SARS-CoV-2 genome, antigens from the Spike protein (S) exhibited the highest rate (26.6%) of cell capture (SCTs that captured at least 1 cell) (Fig. 2b, Supplementary Table 2.1). We identified immunodominant epitopes against SARS-CoV- 2 and common viruses across patients, confirming some previously reported epitopes while discovering new ones (Supplementary Fig. 2b). We also found agreement for dominant TCR gene usage for top epitopes in our dataset with reported literature (Supplementary Table 2.3). For example, HLA-A*02:01/S269- YLQPRTFLL is predominately recognized by TRAV12-1 containing TCRs in our dataset, consistent with previous reports^29^. Selected TCRs were validated via in-vitro Lenti-virial transduction and tetramer assay to confirm antigen specificity (Supplementary Table 2.4, Supplementary Fig. 3a, b).

**Fig. 2:**
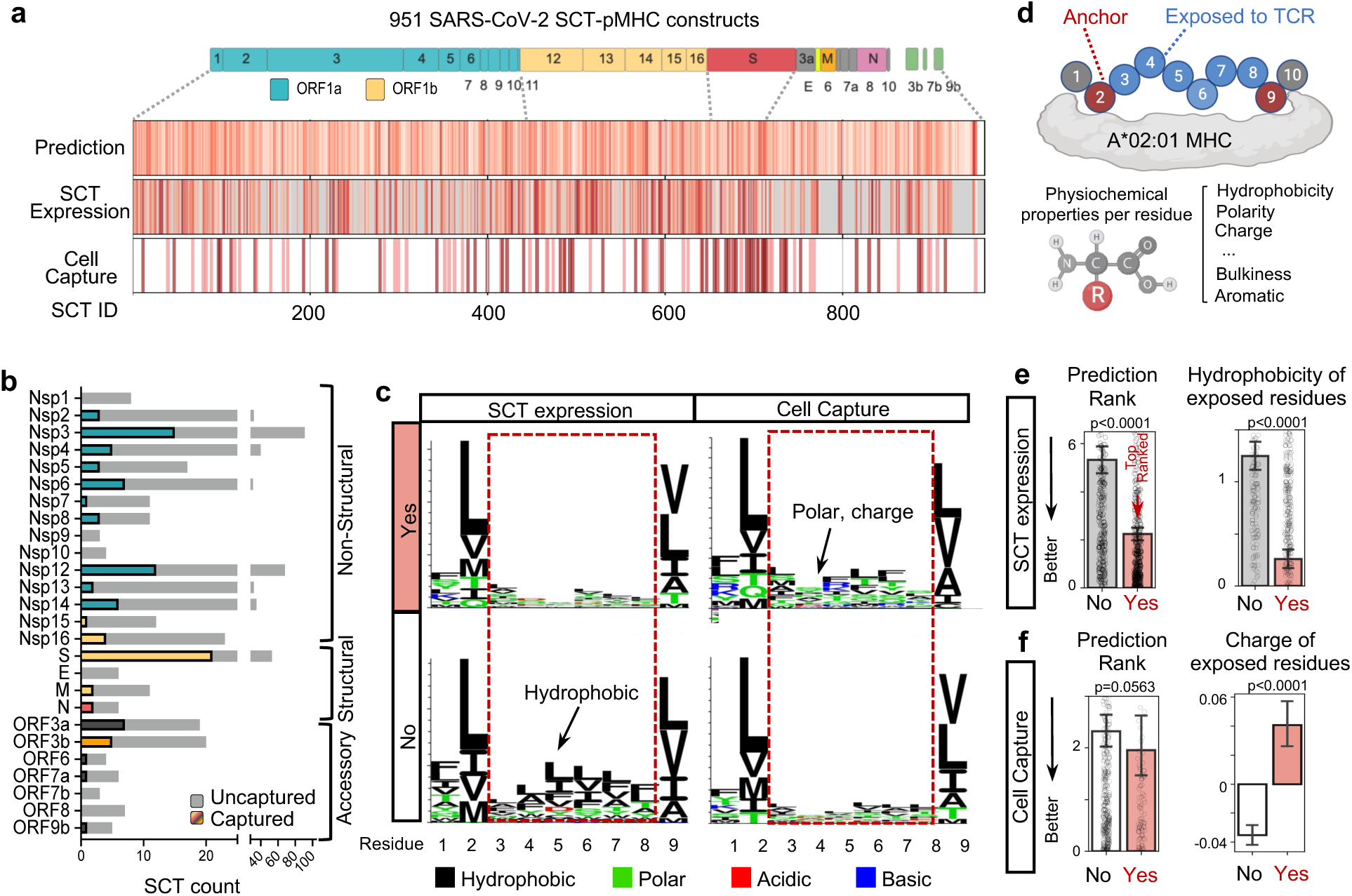
APMAT enables high-throughput representation of whole SARS-CoV-2 genome a. Graphic relating the SARS-CoV-2 proteome to putative HLA A*02:01 restricted antigens (top row), to those SCT constructs that were expressed in usable yield (middle row), and those that captured antigen-specific CD8 T cells from COVID-19 participants (bottom row). For the SCT Expression row, darker red lines correspond to higher SCT expression, while grey means low/no expression. For the Cell Capture row, darker red lines mean more cells captured. b. Bar plot showing the distribution of expressed SCTs for each SARS-COV-2 protein. SCTs that successfully captured CD8 T cells are shown in colored bars. The color code of the captured cells is that used in SARS-CoV-2 proteome of panel a. c. Antigen sequence motif of expressed (top left) vs non-expressed (bottom left) SCT constructs; and those expressed SCTs that did (top right) or did not capture CD8 T cells (bottom right). d. For each peptide, conventional anchor positions (R2, R9) and non-anchor residues are assigned. The physicochemical properties are tabulated for each residue. e. Bar plots comparing expressed (N = 560) vs non-expressed (N = 391) SCT constructs. X-axis: SCT expression status. Note that two of the expressed SCT constructs were not included for 10X experiment due to low sample volume. Y-axis: NetMHC prediction rank (left) and average hydrophobicity of exposed residues (right). f. Bar plots comparing expressed SCTs that captured CD8 T cells (N = 102) vs those that did not (N = 456). X-axis: Cell capture status. Y-axis: NetMHC prediction rank (left) and average charge of exposed residues (right). Charge values represent the average of positive and negative charges rather than the absolute value. For bar plots, data are presented as mean values +/- SEM, with corresponding individual data points overlayed as hollow dots when possible. Dots outside of the range of y-axis are not shown. The Statistical significance was determined using the two-sided Mann-Whitney U test, and p values are annotated on all relevant plots with exact p-values provided unless p < 0.0001.

We next investigated the antigen sequences associated with SCT expression and cell capturing. In Fig 2c (left) we provide a sequence logo representation of the 9-mer SCTs that were or were not expressed, and those expressed as SCTs that did or did not capture cells (Fig. 2c right). We observed an enrichment of hydrophobic amino acids in non-anchor residues for non-expressed SCT constructs (Fig 2c left bottom). For SCTs that captured cells, polar and charged residues such as Threonine (T) and Arginine (R) were enriched in non-anchor positions (Fig 2c right top). We next generated a matrix for each peptide that contained the NetMHCpan prediction, the amino acid identities, and the numeric physicochemical properties (such as hydrophobicity, polarity, etc) for peptide anchor residues (Anchor) and residues exposed to the TCR (Exposed) (Fig. 2d, Supplementary Table 3). As expected, expressed SCTs showed better NetMHCpan prediction (lower prediction rank and lower predicted binding affinity) relative to non-expressed SCT constructs (Fig. 2e left, Supplementary Fig. 4a). Notably, SCT expression yield did not correlate with predicted affinity (Supplementary Fig. 4b). However, the physicochemical properties analysis allowed for a quantitative validation of the hydrophobic trends of non-expressed SCTs (Fig 2e right), and the polar/charged residue trends of cell-capturing SCTs (Fig 2f right). We also show the agreement between SCT expression and NetMHCPan prediction. We categorized peptides as weak-binders, medium-binders, or strong-binders based on NetMHCPan prediction against the A*02:01 MHC molecule (See Methods). In fact, our observations that peptides that are expressed as an SCT exhibit a lower average hydrophobicity closely mirror the NetMHCPan pan predictions. The strong-binders indeed exhibit a relatively higher polarity and a lower hydrophobicity relative to all attempted A*02:01 SCT constructs (Supplementary Fig.4c). In addition, for those strong binders, the hydrophobicity and polarity of expressed vs non-expressed SCTs are not significantly different (Supplementary Fig.4d). Finally, we found that the likelihood that an SCT would be successfully expressed strongly agrees with prediction. For weak-binders, SCT expression rate is 42.0%; For strong-binders, SCT expression is elevated to 82.5% (Supplementary Fig.4e). Hence, the potential SCT- expression biases match closely with the NetMHCPan predictions with respect to the physicochemical properties of the putative epitopes, suggesting that little or no bias originates from the SCT expression itself.

Overall, APMAT enabled a direct and comprehensive analysis of putative epitopes, supporting prediction algorithms for the common HLA A*02.01 allele. The data further suggests that an analysis of the physicochemical properties of the antigens (and possibly their cognate TCRs) may provide insights for interpreting the large multimodal data set generated through APMAT.

### Three peptide groups distinguished by sequence physicochemical properties

We probed for potential relationships between the physicochemical properties of putative epitopes and the antigenicity of those epitopes. We encoded the 951 peptides by residue-level descriptors of their physicochemical properties (Fig. 3a, Methods). Unsupervised clustering based on peptide amino acid identity and properties resulted in a two-dimensional peptide Uniform Manifold Approximation and Projection (Pep-UMAP) (Fig. 3b, Supplementary Table 4). The upper right wing of this UMAP exhibits higher hydrophobicity and bulkiness, and is dominated by non-expressed SCTs. Conversely, the lower left wing displays greater polarity, and is enriched for expressed SCTs (Fig. 3c, Supplementary Fig. 5a). However, SCTs that successfully captured T cells are uniformly distributed across the UMAP, indicating that additional factors beyond hydrophobicity modulate antigenicity.

**Fig. 3:**
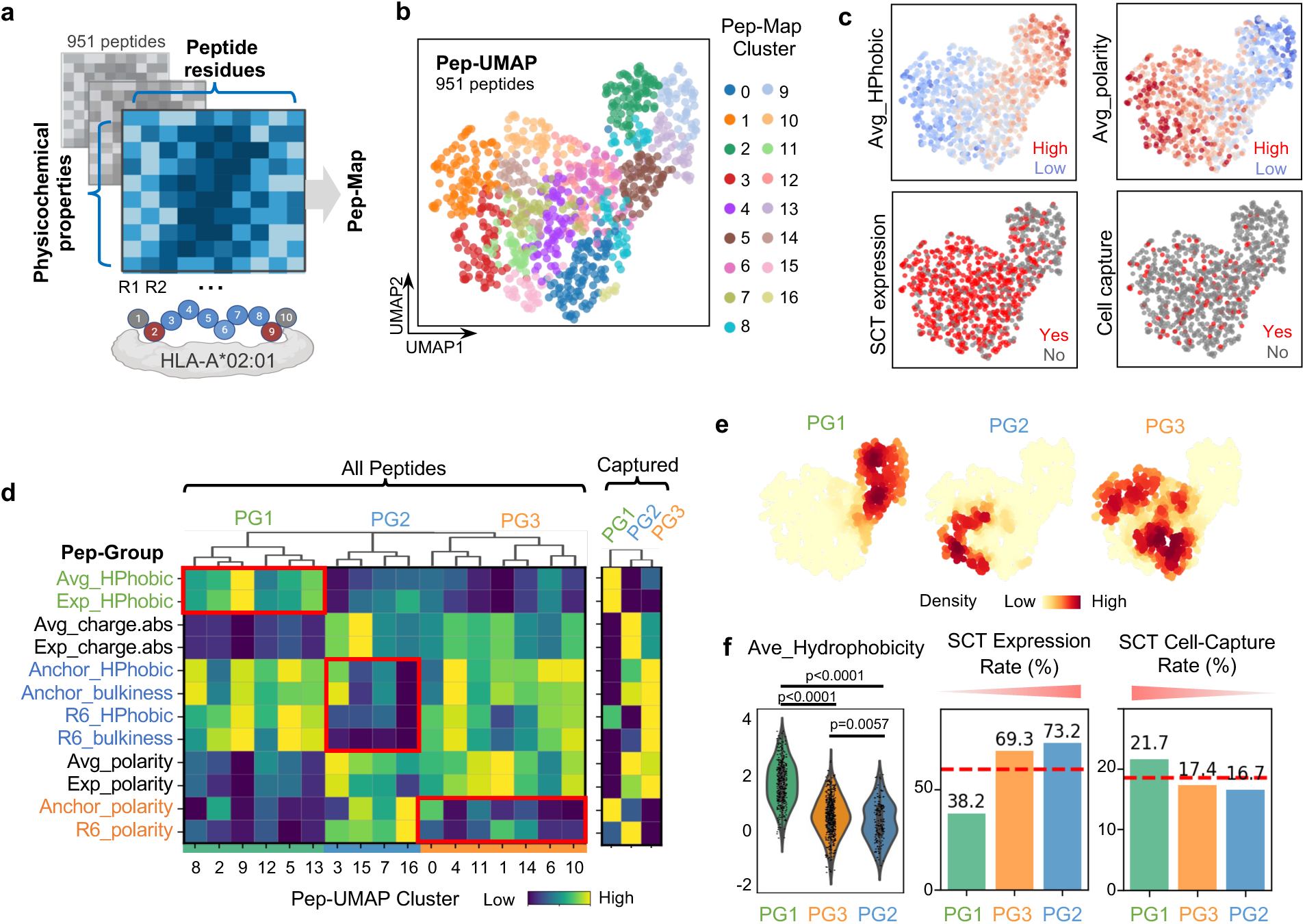
Three peptide groups distinguished by sequence physicochemical properties a. All 951 putative SARS-CoV-2 peptides for HLA-A*02:01 are encoded by the physicochemical properties of amino acids at each position for unsupervised clustering. b. UMAP embedding of all peptides (Pep-UMAP) based on their physicochemical properties with peptide clusters colored (legend on the right), each dot represents a unique SARS-CoV-2 peptide. c. Pep-UMAP colored with selected physicochemical properties, including average (Avg) hydrophobicity (HPhobic), average polarity, whether the SCT expressed, and whether the SCT captured CD8 T cells. Average values were calculated for all residues, including anchor and exposed residues. d. Left: Clustermap of the 951 peptide antigens by their normalized physicochemical properties, revealing 3 major peptide groups (Pep-Groups). The key signatures that distinguish the individual Pep-Groups are highlighted in red boxes. Right: Clustermap of the Pep-Groups including only those peptides that captured T cells. e. Pep-UMAP with densities of PG1-3 depicted, legend on the bottom. f. Left: Violin plot of peptide hydrophobicity for Pep-Groups, sorted by mean value. Middle: SCT protein expression efficiency for the Pep-Groups. Right: SCT cell capture efficiency for Pep-Groups. Mean values +/- SEM are utilized for violin plots. The Statistical significance was determined using the two-sided Mann-Whitney U test, and p values are annotated on all relevant plots with exact p-values provided unless p < 0.0001.

We next utilized unsupervised clustering to resolve whether combinations of physicochemical properties could be used to further classify the peptide antigens. Such analysis clearly distinguished three peptide groups (Pep-Groups), PG1-3 (Fig. 3d). Specifically, PG1 exhibited higher hydrophobicity yet lower charge and polarity (Fig. 3d left top). PG2 and PG3 both showed higher polarity, but differ in anchor and secondary anchor (Position 6) properties (Fig. 3d left bottom). When analyzing only the cell-capturing SCTs, the hydrophobicity of PG1 becomes even more prominent while PG2-3 distinctions diminish (Fig. 3d right). Accordingly, Pep-Groups occupied different regions on the Pep- UMAP (Fig. 3e, Supplementary Fig. 5b). Comparisons of PG1-3 for hydrophobicity, SCT expression rate, and cell-capture rate (Fig. 3f) reveal the most hydrophobic group (PG1) is the most challenging to express, but also exhibits the highest rate for cell capture. Thus, we categorized all putative antigen peptides into unique Pep-Groups for downstream analysis.

### Pep-groups associate with different T cell phenotypes

We next investigated whether the PG1-3 peptide groups associate with distinct CD8 T cell phenotypes during the acute COVID-19 response. The SARS-CoV-2 SCT-dextramer-positive CD8 T cells were filtered to only include those with an assigned patient ID, antigen specificity, and paired TCRα/β sequences. These cells were then projected onto a gene expression UMAP (GEX-UMAP) based on the scRNAseq data (Fig. 4a). Unbiased clustering and differential gene expression analysis defined canonical CD8 T cell phenotypes including naïve, central memory (CM), effector memory (EM), hybrid, and cytotoxic phenotypes (Fig. 4b). For instance, CCR7, LEF1, TCF7, and SELL were up- regulated in naïve and central memory (CM) cells, while memory markers such as IL7R, GZMK were elevated in CM and effector memory (EM) phenotypes (Fig. 4b, Supplementary Fig. 6a-b). Phenotype assignment based on transcriptomics was further validated by surface protein expression measured by scCITE-seq (see Methods) (Supplementary Fig. 6c). As expected, cytotoxic and EM phenotypes dominate the SARS-Cov-2-specific CD8 T cell response during acute disease^26,30,31^, with elevated effector markers such as GZMB, GZMA and PRF1. In addition to gene expression, in Fig. 4c we projected the polarity of exposed residues on the antigen recognized by the individual T cells (left), the distribution of T cells captured by a given antigen across multiple participants (middle), and all captured SARS-CoV-2 specific T cells for a given participant.

**Fig. 4:**
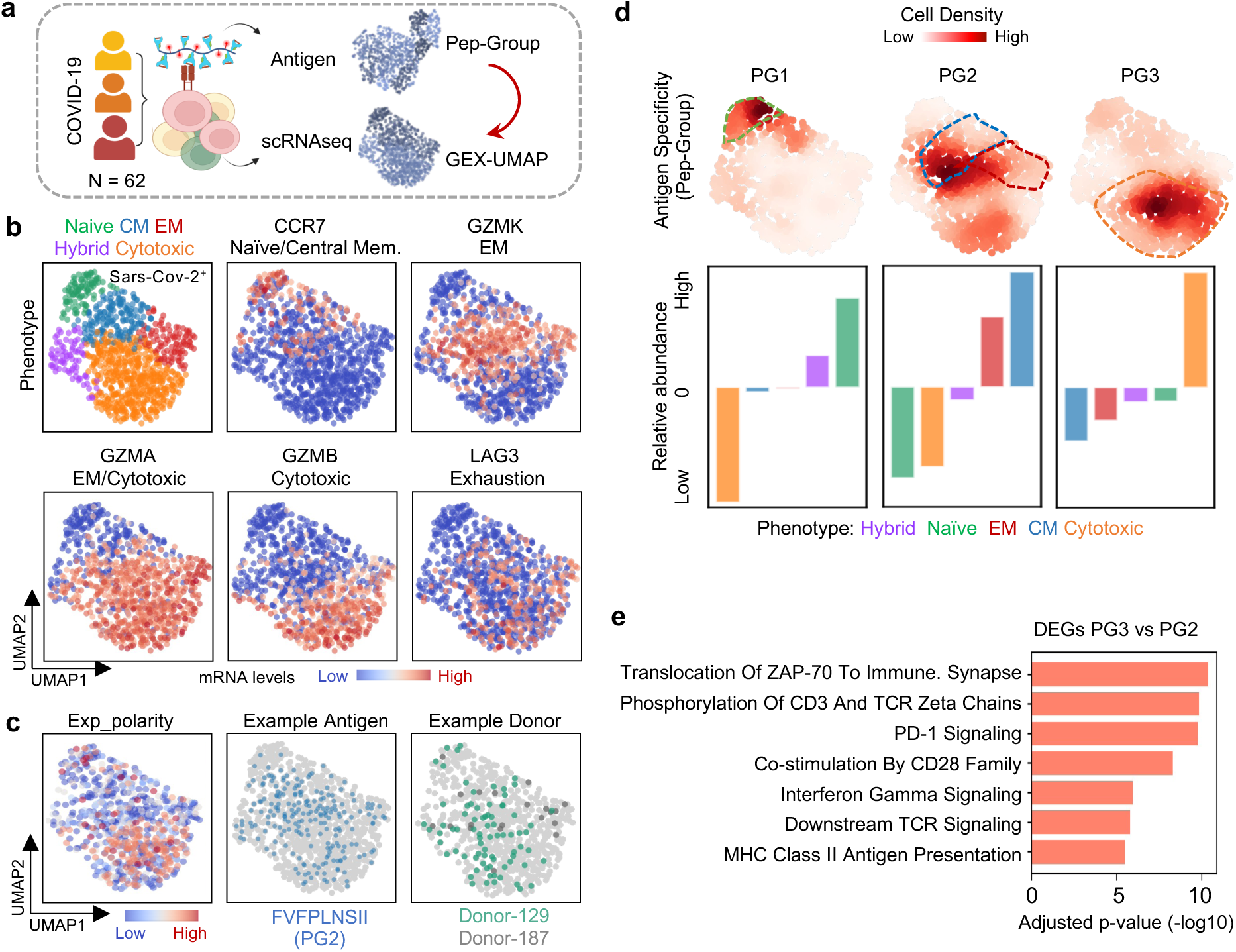
Pep-groups associate with different T cell phenotypes. a. Illustration of the mapping of peptide physicochemical properties on to T cell gene expression profiles. CD8 T cells were captured by SCT-dextramers from 62 HLA-matched COVID-19 participants for scRNAseq and TCR sequencing. Gene expression UMAP (GEX-UMAP) was generated based on scRNAseq. We used the SCT identity to connect Pep-Groups with gene expression. b. GEX-UMAP of Sars-CoV-2-specific CD8 T cells with different phenotypes color-encoded (legend on the top), and then color-coded by expression levels of selected mRNA transcripts. c. GEX-UMAP, color-coded by (left) the polarity of exposed peptide antigen residues; (middle) all T cells specific to a given antigen; and (right) SARS-CoV-2 specific T cells from two study participants. d. Top: GEX-UMAP color encoded with T cell densities specific for antigens from each Pep-Group. Bottom: Bar plot of the relative abundance of phenotypes for T cells associated with each Pep-Group. Note that the UMAP densities in the original were calculated as an odds ratio. e. Top enriched biological pathways of genes significantly elevated in cells captured by PG3 relative to PG2. Adjusted p-values are generated by EnrichR

The projection of antigen polarity on the GEX-UMAP (Fig. 4c) showed a strong skewing towards cytotoxic CD8 T cell phenotypes, suggesting that antigens with particular physicochemical properties may associate with specific T cell phenotypes. We explored this concept by projecting each cell’s antigen specificity (Pep-Groups) onto the GEX- UMAP (Fig. 4d top). PG1-specific cells were dominated by naïve-like phenotypes and exhibited less clonal expansion relative to the other two groups (Supplementary Fig. 6d). PG2-specific cells were enriched with EM and CM phenotypes, and PG3-specific cells were mainly cytotoxic (Fig. 4d bottom). Furthermore, differentially expressed genes (DEGs) in PG3-captured cells have enriched pathway signatures related to immune synapse formation, PD-1 signaling, CD28 co-stimulation, which further highlighted the effector state of PG3-specific cells relative to PG2 (Fig. 4e, Supplementary Table 5-6). This analysis suggests strong associations exist between the physicochemical properties of the peptide antigens, and the phenotypic characteristics of the T cells specific to those antigens.

### TCR hydrophobicity is an important factor for effector function

Intrigued by the link between antigenic peptides and T cell phenotypes, we investigated whether a similar connection exists for the physicochemical features of each antigen-associated TCR-CDR3 sequence by overlaying those features on the GEX-UMAP (Fig 5a, Supplementary Fig. 7a). Specifically, we categorized CDR3β residues into V, J, and CDR3βmer (central) regions: The V and J regions comprise the highly conserved n- and c-terminal motifs respectively; while the central CDR3βmer region, which primarily contacts the antigen, is the most diverse in length and amino acid usage. As expected, cytotoxic cells showed maximal clonal expansion followed by effector memory (EM) cells (Supplementary Fig. 7b). We compared the CDR3β features of effector cells (cytotoxic and EM) relative to other cell types (Supplementary Table 7-8). TCR feature differences were not significant for the V and J regions as expected (Fig. 5c middle). However, for the CDR3βmer region, a preference of sequence features was observed. Effector T cells were marked by higher CDR3βmer hydrophobicity and bulkiness, lower polarity and charge, and shorter length (Fig. 5c top and bottom). Note that CDR3β hydrophobicity was defined both by the percentage of hydrophobic residues, and by the average hydrophobicity across central residues (See method). We validated the above trends by plotting how the selected CDR3β physicochemical properties varied across all cell phenotypes. The CDR3β sequences displayed the trend of increased hydrophobicity and decrease in charge and length from naïve, to EM and cytotoxic phenotypes (Fig. 5d).

**Fig. 5:**
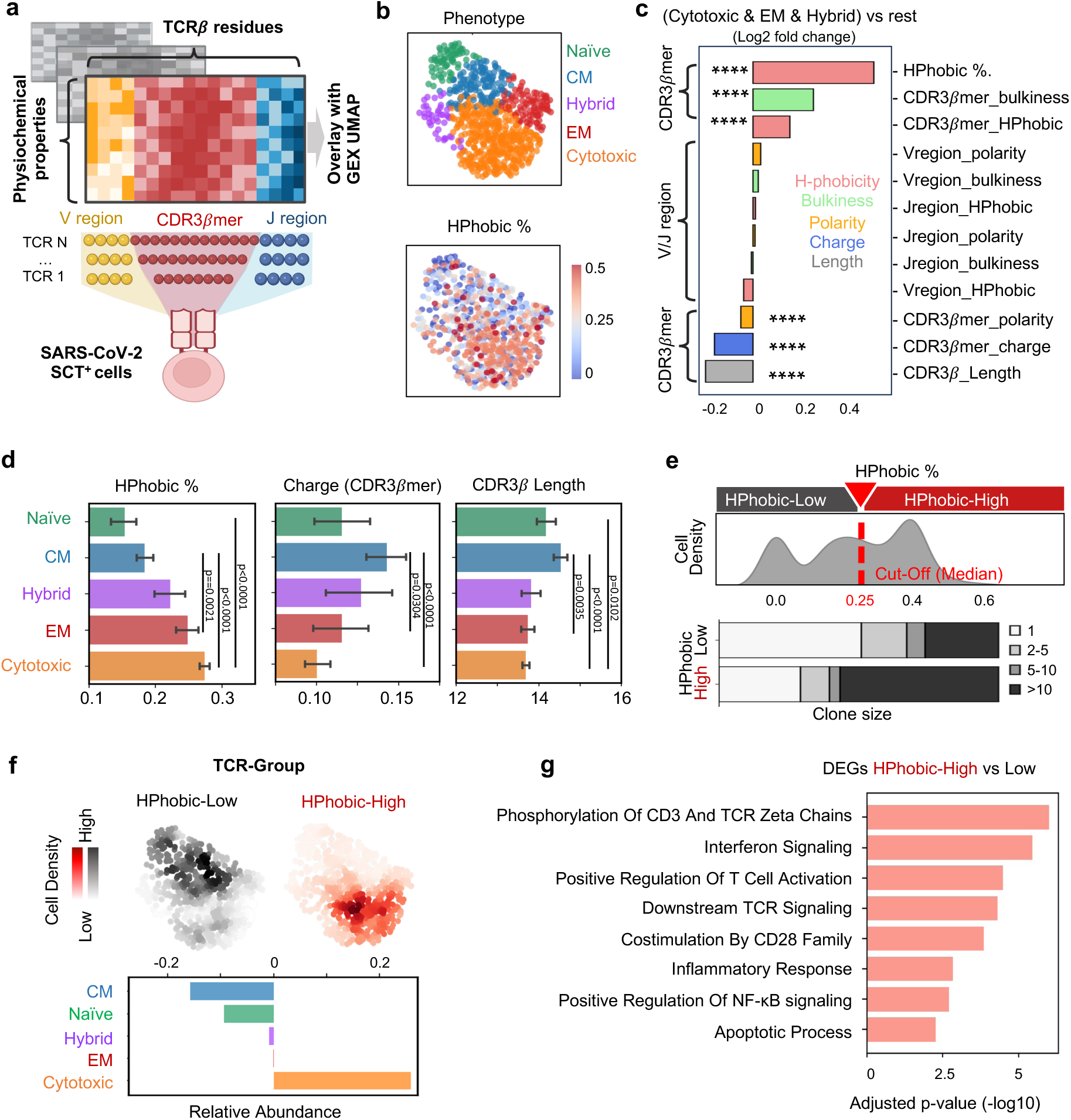
TCR hydrophobicity is an important factor for effector function. a. Each TCR beta chain was split into V, CDR3βmer, and J regions, and then encoded by the physicochemical properties of amino acids at each residue position for overlay on the GEX-UMAP. b. GEX-UMAP, color-coded by phenotypes (top) and overlayed with the percentage of hydrophobic residues within CDR3βmer (HPhobic %) (bottom). c. The top differential TCR physicochemical properties for effector (Cytotoxic, EM and Hybrid) phenotypes, relative to non-effector (CM and naïve) phenotypes. The Statistical significance was determined using the two-sided Mann-Whitney U test, p-values < 0.0001 were marked with ****. See color key inset. Source data are provided as a Source Data file. d. Bar plots showing the variation of selected TCR physicochemical properties across T cell phenotypes. Number of cells: Naïve (n = 68), CM (n = 133), Hybrid (n = 43), EM (n = 93), Cytotoxic (n = 384). For bar plots, data are presented as mean values +/- SEM. p values are annotated on all relevant plots with exact p-values provided unless p < 0.0001. Source data are provided as a Source Data file. e. Top: Binary TCR-Groups (HPhobic-Low and HPhobic-High) are defined based on the median value of hydrophobic residue percentages for all SARS-CoV-2 cells. Bottom: Stacked bar plot of clonal size distribution for each TCR-Groups with legend on the right. f. GEX-UMAP with densities of the TCR-Groups projected. The relative abundance of phenotypes in HPhobic-High cells is plotted at bottom. The UMAP densities were calculated as an odds ratio. g. Top enriched biological pathways of genes significantly elevated in cells with HPhobic- High vs HPhobic-Low TCRs. Adjusted p-values are generated by EnrichR.

The hydrophobicity of the CDR3β exhibited the strongest significant association with T cell phenotype. This prompted us to define a binary classifier (HPhobic-High and HPhobic-low) based on the percentage of hydrophobic residues in CDR3βmer (cutoff = 25%) (Fig. 5e top, see Methods). Cells expressing HPhobic-High TCRs were more clonally expanded than HPhobic-Low ones (Fig. 5e bottom). We first validated that TCR- Groups still preserve the physicochemical features observed earlier: Indeed, HPhobic- High TCRs exhibited higher hydrophobicity, shorter CDR3β length, and lower charge (Supplementary Fig. 7c). Density mapping validated that HPhobic-High TCRs were more prevalent in cytotoxic cells than in memory and naïve subsets (Fig. 5f), with elevated exhaustion markers such as LAG3 and TIGIT (Supplementary Fig. 7d). Gene set enrichment analysis linked HPhobic-High clonotypes to TCR activation (e.g. CD3 and TCR Zeta-chain phosphorylation), inflammation, and apoptosis pathways (Fig. 5g, Supplementary Table 9-10). To test whether dominant clonotypes affect our result, additional analysis was performed by removing dominant TCRs, although dominant TCRs were only found for a few of the epitopes in our dataset (Supplementary Fig. 8a, Details in Method). We observed consistent results with or without the dominant clones. Cells with hydrophobic-high CDR3βs consistently showed elevated PRF1 and GZMB gene expression, as well as higher cytotoxic scores relative to cells with hydrophobic-low CDR3βs after removal of large clones (Supplementary Fig. 8b top). On contrast, hydrophobic-low CDR3βs associated with higher expression of naïve and memory related genes including CCR7 and IL7R (Supplementary Fig. 8b bottom). These results indicate that our original conclusions are not biased by specific dominant clones.

Overall, we demonstrated that, the physicochemical properties of both the peptide antigen and the TCR CDR3β exhibit strong associations with T cell phenotype for SARS-CoV-2 specific CD8 T cells.

### Integrated analysis of both peptide and TCR physicochemical features orchestrate phenotypes of SARS-CoV-2-specific CD8 T cells in acute disease

To elucidate how interplay between antigen and TCR interactions influence T cell function, we systematically linked peptide and TCR features for the subset of SARS-CoV-2 CD8 T cells with fully paired antigen-TCR information (See Method). This encompassed 87 unique antigenic peptides (SCT-pMHCs) and 440 paired TCR clones (Fig. 6a). Notably, distinct TCR Groups (HPhobic-Low and HPhobic-High) were discerned within each Peptide Group, prompting a refined categorization into PG-TCR Groups based on combined antigen peptide and TCR features (Fig. 6b). For example, PG3-High denotes cells captured by PG3 peptides with HPhobic-High TCRs.

**Fig. 6:**
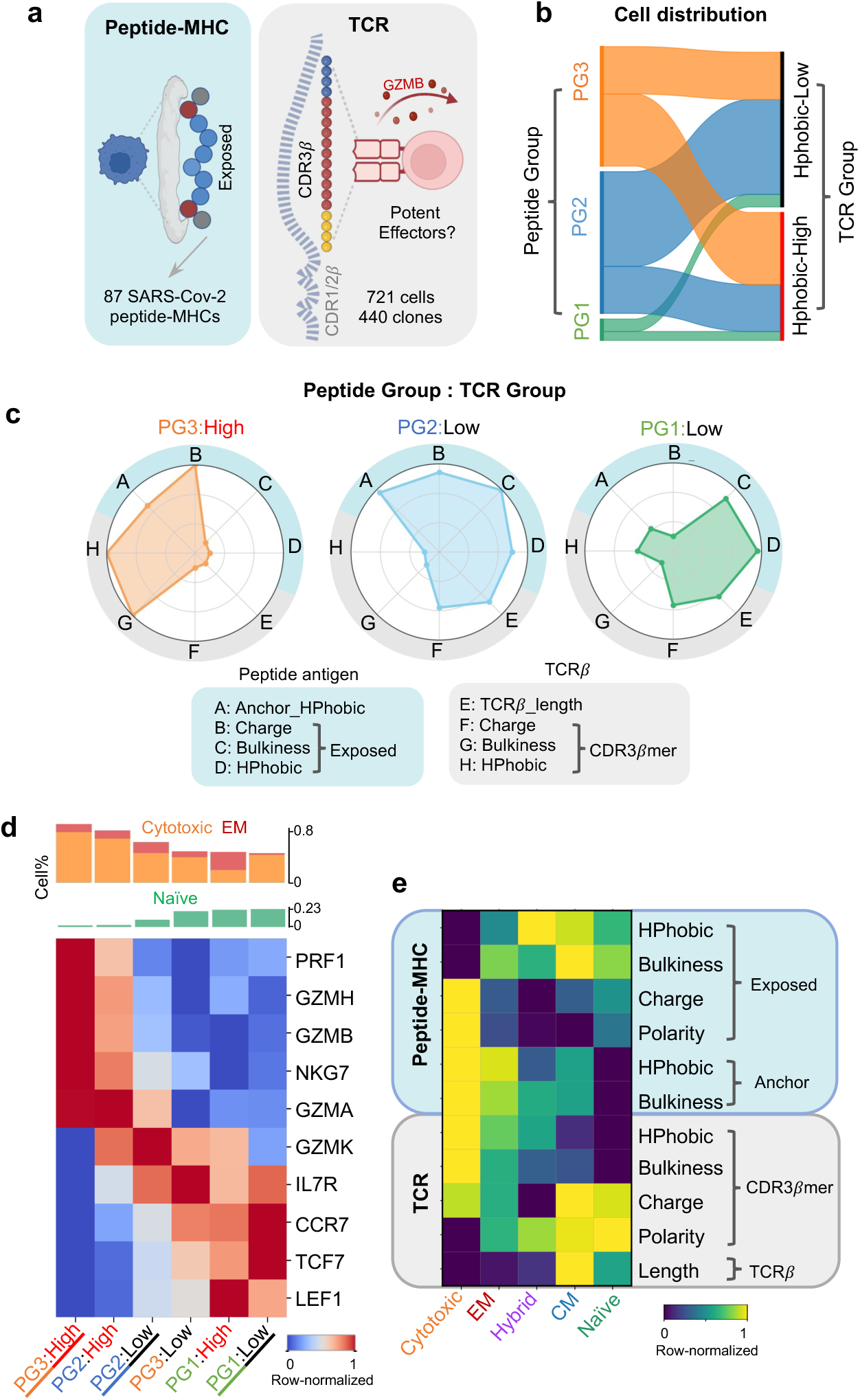
Combination of peptide-TCR features associated with cell phenotypes a. We investigated the combination of peptide and TCR properties for antigen-TCR paired SARS-CoV-2 CD8 T cells. The plotted peptides are those that captured cells. b. Cell distribution between peptide groups (Left) and TCR groups. c. Radar graphs of physicochemical properties for representative PG-TCR groups. Blue and grey shaded areas of the outer rings indicate Pep-Group, and TCRβ properties, respectively. Each axis displays the normalized average value for each property, with lowest value in the center. The shaded polygons reflect the property space occupied by the peptide-TCRβ groupings. Legend on the bottom. d. Top: Cell percentage of Cytotoxic, EM and Naïve phenotypes for each PG-TCR group. Bottom: Heatmap showing selected mRNA levels for each PG-TCR group. Underlined PG-TCR groups are those from panel c. e. Heatmap showing average value of peptide and TCR physicochemical properties for each phenotype.

We then further evaluated the combinatorial effect of peptide and TCR features for each PG-TCR group. We depicted key physicochemical properties identified earlier on radar plots, revealing a significant shift in overall characteristics between PG-TCR groups (Fig. 6c, Supplementary Fig. 9a). For example, PG3:High exhibits high peptide charge and low hydrophobicity on exposed residues, combined with high CDR3βmer hydrophobicity and bulkiness (Fig. 6c left). In contrast, PG2:Low (Fig. 6c middle) emphasizes CDR3βmer length and charge, while PG1:Low (Fig. 6c right) emphasizes distinct characteristics compared to PG3:High.

Building on our findings, we explored how PG-TCR groups associate with T cell phenotypes. PG3:High cells displayed the strongest cytotoxicity – marked by the highest percentage of effector cells and elevated cytotoxic cytokines (e.g. GZMB, PRF1, GZMH) (Fig. 6d left, Supplementary Fig. 9b-c). PG1:Low describes cells with TCR:pMHC interfacial properties that are opposite that of PG3:High, and those cells similarly exhibit phenotypes that contrast with PG3:High – marked by highest frequency of naïve cells and elevated naïve-associated markers (e.g. CCR7, LEF1, TCF7) (Fig. 6d right). Notably, PG1 consistently exhibits a Naïve-like phenotype, regardless of TCR-group. With intermediate peptide-TCR properties, PG2:Low and PG3:Low represent transitional phenotypes. These associations between PG-TCR groups and gene expression were validated using protein markers by integrating existing scCITE-seq data (See Methods) (Supplementary Fig. 9d). We performed further analysis by directly linking the key physicochemical properties to phenotypes independent of the PG-TCR groups. This heatmap highlights the balance between the properties of the antigen and the TCR CDR3b sequence (Fig. 6e), and illustrates the near opposite relationships between naïve/CM cells relative to cytotoxic cells at the stage of acute disease. Hence, by integrating peptide and TCR sequence features, APMAT reveals fundamental rules in the peptide-TCR-phenotype synergy.

### Distinct peptide-TCR groups associate with distinct longitudinal fates of SARS-CoV-2-Specific CD8 T cells

T cell activation strength through pMHC-TCR interaction can influence cell fate over time^32–34^. Certain T cell phenotypes exhibit long-term *in vivo* persistence following acute illness, while highly cytotoxic phenotypes can undergo activation-induced cell death (AICD) and contract after clearance of pathogen^35–37^. We hypothesized that distinct Pep- TCR groups may associate with T cell fate decisions across longitudinal disease trajectories as well. Leveraging the longitudinal scRNA-seq and scCITE-seq data generated from the same COVID-19 cohort^25^, we tracked SARS-CoV-2-specific CD8 T cells matched on patient ID and TCR sequences from acute to post-acute timepoints (Fig. 7a, Supplementary Table 11). This analysis further revealed longitudinal differences between PG-TCR groups. As expected, the overall percentage of SARS-CoV-2 specific CD8 T cells identified from the reference dataset decreased from 3% to 1.3% at the post-acute timepoint. Specifically, PG3:High cells were short-lived, showing the greatest drop in abundance at the post-acute timepoint. By contrast, PG1 cells (including PG1:High and PG1:Low) were the most persistent, showing an increased abundance at the later timepoint (Fig. 7b, Supplementary Fig. 9e).

**Fig. 7:**
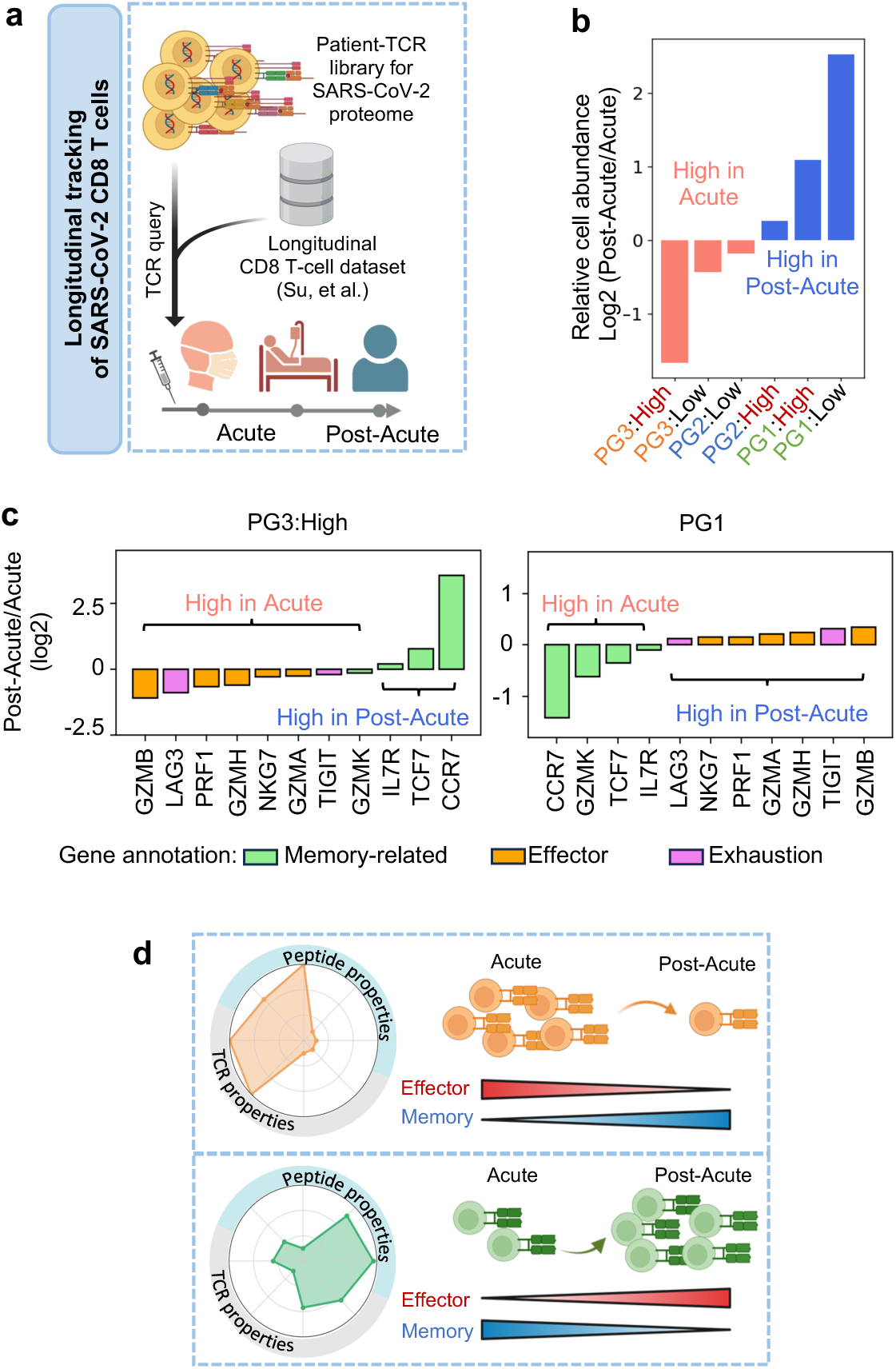
Longitudinal analysis for PG-TCR groups a. Tracking SARS-CoV-2 CD8 T cells from acute to post-acute timepoint based on TCR- GLIPH query from a previously reported longitudinal dataset. b. Cell abundancy changes from post-acute to acute timepoint for each PG-TCR grouping. Red or blue means higher abundancy at acute or post-acute timepoint, respectively. c. Selected gene expression changes from the acute to post-acute timepoints for each PG-TCR groups. Gene annotations on the bottom. d. Summary: The association between distinct peptide-TCR properties and cell fates for antigen-specific CD8 T cells.

In addition to abundance changes over time, we examined gene expression changes for SARS-CoV-2 specific CD8 T cells. Combining all PG-TCR groups, we observed that persisting cells at the post-acute timepoint showed higher expression of genes that were associated with long-lived memory signatures (such as CCR7, IL7R, HLA-DR, MKI67, etc) (Supplementary Fig. 9f), in agreement with previous studies^16,18,20^. In addition, our analysis further suggested a few other trends. Specifically, PG1 cells identified at the late timepoint showed a relatively lower naïve signature (CCR7, TCF7, etc) and slightly higher effector functions (GZMB, GZMH, etc) than those cells at the earlier timepoint (Fig. 7c right, Supplementary Fig. 9g right). By contrast, the few remaining post-acute PG3:High cells evolve to express lower cytotoxicity (GZMB, PRF1, etc) and higher CCR7, indicating a shift towards less effector or central memory phenotypes^16,38,39^ (Fig. 7c left, Supplementary Fig. 9g left). In summary, APMAT revealed that distinct combinations of peptide and TCR physicochemical properties exhibit clear associations with not only cellular phenotypes at acute disease, but also divergent cell fates over time (Fig. 7d).

## Discussion

The conceptual advance we report as APMAT lies in the combination of the large and comprehensive experimental data set and the biophysical analysis of the TCR-pMHC interface. APMAT provides a framework for the integrated analysis of antigen-specific CD8 T cells paired with phenotypic data on those T cells. We applied APMAT to investigate circulating CD8 T cells collected from 62 participants at the acute and convalescent stages of COVID-19. These cells exhibited specificity to SARS-CoV-2 antigens presented by HLA A*02:01 from across the full viral proteome. Our analysis uncovered relationships between the physicochemical characteristics of the pMHC:TCR interface and the corresponding T cell phenotypes and T cell phenotypic evolution. Our analysis is, in several aspects, aligned with existing literature. We identified T cell populations targeting previously reported immunogenic antigens^17,24,40,41^, and we find that the prediction rankings from the commonly-used NetMHCpan algorithm effectively assist in designing pMHC multimer constructs that will most likely capture T cells. In addition, the longitudinal behavior of the antigen-specific clonotypes is consistent with current literature. Highly-cytotoxic effector clonotypes, which are expanded at acute disease, contract significantly at the time of convalescence, potentially from antigen-induced cell death. In contrast, those persisting clonotypes with mild effector properties evolve towards central or effector memory phenotypes^42^. Further, clonotypes that exhibit memory and progenitor-like phenotypes at acute disease persist or expand at convalescence^16,43,44^.

APMAT analysis uncovers novel relationships between viral antigen-specific T cells, TCR clonotype, and T cell phenotype for the common HLA allele A*02.01. Take, for example, the above-described case of effector T cells that expand during an acute infection and contract at convalescence^42^. Our analysis further suggests that T cells possessing TCRs characterized by a hydrophobic CDR3β, which recognize antigens featuring hydrophobic anchor residues alongside charged or polar exposed residues, are statistically more inclined towards effector phenotypes at the stage of acute COVID-19, and contract during convalescence. An analogous description of the pMHC:TCR interface (but with near opposite characteristics) can be used to identify those clonotypes that exhibit naïve or central memory phenotypes at acute disease and remain at convalescence. We focus on CDR3β, but CDR3α chains can be similarly analyzed to identify specific physicochemical properties that significantly associate with effector (Cytotoxic, EM, Hybrid) or non-effector T cells. Those TCRα properties are distinct from those for TCRβ, suggesting that the influences from the α and β chains might be complementary rather than independent (Supplementary Fig. 10a).

An additional significant aspect of this study is the detailed characterization of paired peptide antigen with TCR. Notably, our analysis reveals that the anchor and exposed residues of the antigen exert different influences on T cell phenotype. Unlike many prior studies that relied on a rough annotation of the antigen identity, our approach takes into account the position of each individual residue, as well as the classification of antigens that may be comprised of distinct sequences, and yet exhibit biochemical similarities at the TCR:pMHC interface.

Whether or not these physicochemical determinants are general to other disease contexts or other HLA Class I alleles is not resolved here, although the consistency of the picture painted here with what has been observed in tissue settings in murine models of chronic viral infection,^4^ as well as in tumors,^5,6^ does suggest some level of generality.

Further, an analysis of a separate study on SARS-CoV-2 reactive T cells (Supplementary Fig. 11a)^45^, as well as our recently reported phenotypic analysis of T cell clonotypes specific to three highly immunogenic viral antigens, including two from influenza and CMV^8^, (Supplementary Fig. 11b) also suggest a degree of generality. Datasets of similar breadth and depth for a range of diseases and for antigens presented by different HLA alleles are needed to more fully resolve this picture, and such work represents an exciting future direction.

We hypothesize that an analogous study as the one reported here, but directed at antigens presented by different HLA alleles, might yield a different set of rules that are dependent upon HLA-specific docking geometries. While Class I HLA A, B, and C alleles are highly polymorphic, over 90% of the world’s population carries at least one of the dozen most common. An APMAT analysis centered around those most common alleles should offer valuable insights for assessing the therapeutic potential of tumor-targeting TCRs and T cell vaccines.

## Methods

### Lead Contact

Further information and requests for resources and reagents should be directed to and will be fulfilled by the Lead Contact, Dr. James R. Heath.

### Participants and sample collection

The study cohort is a subset of the INCOV cohort published previously^25,26^. Procedures for the INCOV study were approved by the Institutional Review Board (IRB) at Providence St. Joseph Health with IRB study number STUDY2020000175 and the Western Institutional Review Board with IRB study number 20170658. This research complies with all relevant ethical regulations. Potential participants were identified at five hospitals of Swedish Medical Center and affiliated clinics located in the Puget Sound region near Seattle, WA. All enrolled participants provided written in-person informed consent and samples were de-identified prior to analysis. 62 HLA A*02:01 individuals from the INCOV cohort were selected for this study. PBMCs collected at Acute timepoint, including enrollment close to diagnosis (Acute-1) and 1 week (Acute-2), were used for antigen-TCR paring assay in this study.

### Large-scale preparation of peptide-HLA complex libraries

Single chain trimer (SCT) peptide-MHC (pMHC) libraries of the virus antigens were generated as described previously^23^1/8/25 4:32:00 PM. Briefly, potential HLA A*02:01 binding epitopes (9-11 mer peptides) were generated from the complete SARS-CoV-2 genome (Wuhan-Hu-1 strain, GenBank ID: MN908947.3) and filtered by NetMHCpan-4.1 prediction^28^. We categorized peptides as weak-binders (column BindLevel = 0), medium-binders (column BindLevel = WB), or strong-binders (column BindLevel = SB) based on NetMHCPan prediction. Note that even the weak-binders are relatively good candidates compared to peptides that were not attempted for SCT expression. A plasmid library of pcDNA3.1 vectors encoding covalently linked peptide antigen, β2M, HLA was built and verified by SANGER sequencing. Plasmids were transfected into Exp293 cells (ExpiFectamine™, Thermo Fisher) following manufacturer protocol. SCT expression yield was measured and normalized. Expressed pMHC-like SCTs were biotinylated (BirA ligase Kit, Avidity) and Histag purified (IMAC PhyTip columns, PhyNexus) in 96-well format. Individual SCT concentration was measured by protein absorbance at 290nm.

### SCT-dextramer generation and cell staining

SCT dextramers were individually DNA barcoded using dCODE Klickmers (dCODE Klickmer, Immudex). Briefly, SCT pMHC monomer was mixed with barcoded dCODE- PE-dextramer at a ratio of 20 ligands per dextran and incubated for at least 1h on ice before adding biotin (100 μM) to block free binding sites. Dextramer cocktails were prepared by mixing 31-65 unique SARS-CoV-2 SCT dextramers and CMV (NLVPMVATV) SCT dextramers freshly before cell staining. PBMCs from 62 participants were thawed for CD8 T cell enrichment (Human CD8 T cell Isolation Kit, Miltenyi Biotec) according to the manufacturer’s protocol then incubated with Human TruStain FcX blocking reagent (422302, BioLegend) for 10 min at 4 °C before wash. Cells were then divided into tubes and processed simultaneously on ice. Each tube of CD8 T cells were stained with a cocktail of dextramers for 25 min on ice in the presence of herring sperm DNA according to the manufacturer’s instructions, with individual dextramer concentration at 1.1 nM. Cells were washed three times before surface antibody staining. BV421 anti-human CD8 Antibody (BioLegend, 344748, clone SK1) and Apotracker™ Green viability dye (Biolegend, 427403) was added into each tube, in addition to one unique TotalSeq-C anti-human hashtag antibody (BioLegend) to identify each tube. Samples were incubated for 30 min on ice and washed 3 times before sorting.

## 10X genomics single cell sequencing

Single, live, CD8, dextramer-positive T cells were sorted into FACS buffer (PBS, 2%FBS, 2mM EDTA and 10mM HEPES) using a BD FACSAriaII cell sorter. Sorted cells were immediately were pelleted, resuspended and loaded into a 10X Chromium reaction for single cell RNA sequencing (scRNA-seq). GEX, VDJ and Surface Protein libraries were generated using Chromium Next GEM Single-Cell 5′ kits v2 (10X Genomics) according to the manufacturer’s protocol. Libraries were sequenced on an Illumina NovaSeq at a read length of 26x90 bp.

### Whole genome sequencing

DNA extraction from whole blood was performed via bead-based enrichment on an automated extraction platform (Qiagen Qiasymphony and/or Promega Maxwell). The resultant extracts were quantified by Nanodrop. WGS library preparation was performed using Illumina DNA Prep kits and the final barcoded libraries were quantified by fluorometer. Libraries were multiplexed and loaded onto an Illumina flow cell for sequencing at 30x or higher coverage on a NovaSeq 6000 instrument. Raw sequencing data was analyzed for sequence variants using the Illumina DRAGEN field-programmable gate array (FPGA) platform. Briefly this platform performs the following automated steps: conversion of raw sequencing image data into demultiplexed fastq files, alignment to the reference human genome (hg19), analysis of single nucleotide variants, indels and copy number/structural variants using variant calling algorithms as well as assessment of sequencing data quality. Analyses with hg38 were computed after a liftover was done using the UCSC browser^46^. WGS information was de-identified.

### Single-cell sequencing data processing

Transcriptome, TCR, surface protein levels and antigen specificity were simultaneously analyzed for each cell. Raw data were processed via Cell Ranger Single-Cell Software Suite (v3.1.0, 10X Genomics) using GRCh38 as a reference. Cells that fit any of the following filters were excluded due to low quality: n-counts <1000 or >10,000, n-genes <250 or >2500, mitochondrial percentage >10%. Gene counts for each cell were normalized by total expression, multiplied by a scale factor of 10,000 and transformed to log scale.

### Single CD8 T cell phenotype assignment

Single cells were assigned phenotypes by clusters determined through the leiden algorithm. Phenotype associated transcripts were acquired from literature as follows^16,26,39,47^. Naïve/memory: LEF1, TCF7, CCR7. Memory: IL7R. Effector Memory: GZMK. Cytotoxic: PRF1, GZMB. Exhaustion: TIGIT, PDCD1. Proliferation: MKI67.

### Transcriptomics data validation via scCITE-seq

To validate the transcriptomics data, we leverage our previously reported scCITE-seq data that simultaneously measured transcriptomic and surface protein levels and TCRα/β from the same single cell. We extracted each cell’s scCITE-seq data by TCR-based inquiry (illustrated in manuscript Fig. 7a, and methods). This leads to single cells with matching transcriptomics and surface protein data (which is the data for plotting main Fig. 7a-c, Supplementary Fig.6c, Supplementary Fig.9d). Our phenotype assignment based on transcriptomics (as the reviewer mentioned for Main Figs. 4 and 5) can be validated by surface protein expression (measured by scCITE-seq). These pairs including CCR7 and CD197-CCR7-CITEseq, IL7R and CD127-IL-7RA-CITEseq. In addition, well-established surface protein markers were supported by our transcriptomics data, such as CD45RO for memory phenotype, CD45RA for naïve phenotype, CD25 for central memory^48^.

### Demultiplexing using genetic variants

We wrote a Snakemake workflow (https://github.com/racng/snakemake-merge-wgs) for processing GVCF files from WGS to generate a multi-sample VCF file of exon variants.

GVCFs are combined for specified genomics region (autosomes and sex chromosomes) using GATK (v4.1.9.0) GenomicsDBImport to generate a GenomicsDB datastore, which is then used by GenotypeGVCFs for joint calling of variants. After removing indels using vcftools (v0.1.16)^49^ and excluding intron variants via bcftools (v1.8)^50^ the remaining exon variants were lifted to GRCh38 (hg38) using CrossMap (v0.5.2)^51^ and filtered again to remove KI27 contigs and duplicated variants. For each 10x library, BAM alignment files from cellranger were filtered for reads from autosomes and sex chromosomes. Using the processed VCF file (929,678 SNPs), single cell variants were extracted from the filtered BAM files via 10x Genomics VarTrix (https://github.com/10XGenomics/vartrix) with the coverage scoring method. To reduce memory usage, single cell variants were kept if both ALT and REF alleles are detected in the dataset. We then used vireo (v0.5.0)^52^ to assign donor identity and doublets based on the processed single-cell and WGS variants. Doublets were removed. Cells that unassigned with donor identity were removed.

### Antigen assignment based on dextramers and hashtags

Raw reads for each Hashtags and Dextramers were normalized. Cells were assigned to their maximally expressed Hashtag. Cells that expressed multiple Hashtags at a high level were removed as potential doublets. Hashtag identities were then used to identify cells’ SCT-dextramer cocktail. For each cell, we calculated the number of unique molecular identifiers (UMIs) for each dextramer, and the percentage of each dextramer. We assigned each cell an antigen only if their UMI count was >25 and the UMIs specific for that dextramer occupied >25% of that cells’ dextramer reads. Antigens were then assigned by the maximally mapped dextramer for each cell. Ambiguous cells that didn’t assigned with any dextramer were removed.

### Peptide physicochemical property assignment and Pep-UMAP

We first transform peptide sequences into a numerical peptide matrix, where each row represents a residue position, and each column represents a feature characterizing amino acids, including amino acid identity, charge, hydrophobicity, weight, bulkiness, polarity, sulfur presence, aromaticity (Supplementary Table 3: Amino acid property scales used for peptide and TCR residues)^12^. The numeric values were scaled to ensure consistency range of values. For quantitative comparison of peptides in Figure 2, we calculated the average value for each property for anchor residues and TCR-exposing residues. Specifically, position 2 and 9 (and occasionally 6) tend to serve as anchors for HLA- A*0201 binding, while other exposed residues potentially contact the TCR (CITE). Logo plot in Figure 2 was generated by Seq2Logo - 2.0^53^ using default settings. In figure 3, to visualize peptide features and similarities, we applied UMAP followed by an autoencoder for dimensional reduction. In detail, since peptides have varying lengths, the residue positions are mapped to a normalized scale of 0 to 100. Each amino acid’s features are replicated across the corresponding positions in the matrix. We then implemented an autoencoder using an MLPRegressor from scikit-learn to reduce the dimensionality of the peptide matrix. Finally, we computed a two dimensional peptide UMAP (Pep-UMAP) using scanpy.tl.umap to visualize peptide features and similarities.

### TCR physicochemical property assignment

TCR-related physicochemical properties were computed for each cell based on its TCR CDR3 sequence of the beta chain. These properties, including charge, hydrophobicity, weight, bulkiness, polarity, sulfur presence, aromaticity, were calculated based on Supplementary Table 3 (Amino acid property scales used for peptide and TCR residues). Charge is absolute value unless specifically indicated. Specifically, we categorized CDR3β residues into V, J, and CDR3bmer regions. The V/J region comprise the first four, and last five amino acids, respectively, while the central CDR3bmer contains the amino acids in between. We evaluated TCR CDR3 hydrophobicity numerically by the average hydrophobicity of the CDR3bmer region. Additionally, we introduced a categorical score called HPhobic%. HPhobic% represents the percentage of strongly hydrophobic residues (A, V, L, I, F, M) in the CDR3β, excluding the first and last four amino acids.

### TCR physicochemical property analysis

We calculated 45 properties for each cell’s TCR sequences, including charge, hydrophobicity, weight, bulkiness, polarity, for three regions of CDR3β (V, J, CDR3βmer), as well as full CDR3α chain. The Mann-Whitney U test is applied to compare the distribution of each property between effector (Cytotoxic, Effector Memory, Hybrid) and non-effector (Naïve and Central Memory) cell phenotypes. Log2 fold change (log2fc) for each property was calculated between the mean values of effector cell types and non-effector cell types. Top selected properties are based on the criteria of log2fc absolute value > 0.05, and p value <0.05. (Supplementary Table 7 and 8). For Supplementary Fig. 8b, Dominant TCRs were removed to test whether our result is biased by large clones. In detail, we performed additional analysis by removing T cell clonotypes and then evaluated the relationship between TCR Groups and phenotypes without these dominant clones. Dominant clones were defined as clone size >= 50 based on 10X VDJ library readout (which counted all cells in the pre-filtered dataset). We also defined phenotype scores as: Cytotoxic score (GZMB, GZMA, GNLY, PRF1) and Naïve-Memory score (TCF7, CCR7, SELL, LEF1, GZMK, IL7R). These were calculated by Scanpy.tl.score to represent the average expression of the given set of genes.

### Peptide and TCR groups density analysis

Densities for peptide groups were projected onto GEX-UMAP by matching the antigen specificity of each single CD8 T cell to the peptide group that antigen peptide belongs to. Embedding density was first calculated via sc.tl.embedding_density then a 5 n-neighbor kNN graph was used to diffuse the values via five iterations to create a whole UMAP score for the density scores, as reported previously^8,54^. TCR group density calculations were implemented this same methodology. The UMAP densities in the original Fig. 4d and Fig. 5f were calculated as an odds ratio. In the area of low cell frequency of certain PG/TCR groups, the density has a low value – but not necessarily zero.

### Differential gene expression and signature analysis

Differentially expressed genes were called via scanpy.tl.rank_genes_groups through the Scanpy package using the Wilcoxon method which implements the Mann-Whitney U test^55^. Differentially expressed genes (DEGs) between peptide groups (PG2 vs PG3) were filtered for p values < 0.05 (Supplementary Table 5). Enriched pathways through filtered DEGs were computed via Enrichr^56^ and provided in Supplementary Table 6. Top enriched pathways in PG3 than PG2 were reported in Fig. 4e. DEGs between TCR groups (HPhobic-High vs HPhobic-Low) were called similarly to generate list of DEGs (Supplementary Table 9) and enriched pathways (Supplementary Table 10). Top enriched pathways in HPhobic-High than HPhobic-Low were reported in Fig. 5g.

### Longitudinal T cell inquiry by GLIPH2 analysis for SARS-CoV-2 specific TCRs

We utilized a reference dataset - our previously reported longitudinal dataset on 209 COVID-19 participants contained both scRNA-seq and scTCR-seq data from the same single cell along with assigned donors^25^. We perform GLIPH2 analysis on SARS-CoV-2 specific TCRs identified in this study that assigned with antigen specificity, and TCRs from the longitudinal dataset with unknown antigen specificity. GLIPH2 was run on http://50.255.35.37:8080/ using the GLIPH2 algorithm^57^, version 1.0 reference for CD8, with all_aa_interchangeable set to YES. Antigen specific CD8 T cells were identified as those that belonged to the same GLIPH group of the SARS-CoV-2 specific TCRs identified in this study. Longitudinal SARS-CoV-2 specific CD8 T cells were then subset from the reference dataset for analysis in Figure 7. Cell count for each PG-TCR groups at Acute and Post-Acute timepoints were provided in Supplementary Table 11.

### Statistical analysis

Microarray like datasets were analyzed using SCANpy and statistical comparisons were generated using scanpy.tl.rank_genes_groups using the wilcoxon method. All correlations were calculated using Pearson correlation. All p values were calculated using Mann-Whitney U test unless otherwise specified. Bar charts were provided with error bars when multiple values were present, and these bars represented standard errors. Bar level represented the mean variable value.

## Data Availability

The single cell sequencing data generated in this study have been deposited in the ArrayExpress database under accession number: E-MTAB-14002, or using the URL: https://www.ebi.ac.uk/biostudies/arrayexpress/studies/E-MTAB-14002?key=3ae4f467-fe01-4d03-b2fe-8a6a522e1cabA.

Source data in this study are provided in the Supplementary Information/Source Data file. Any additional information required to reanalyze the data reported in this work paper is available from the Lead Contact upon request.

## Acknowledgements

We are grateful to all participants in this study and to the medical teams at Providence Swedish Medical Center and UW for their support. We thank the Core Facility at Institute for Systems Biology, the Northwest Genomic Center and Fred Hutchinson Cancer Research Center for help with sequencing services, and the ISB-Swedish COVID- 19 Biobanking Unit. We acknowledge funding support from the Parker Institute for Cancer Immunotherapy (J.R.H., P.D.G.), Merck, the Biomedical Advanced Research and Development Authority (HHSO10201600031C to J.R.H.), the NIH (1 R01 CA264090-01 to J.R.H. and P.D.G, and CTSI UL1TR001881 to A.M.X). Y.S. was supported by the Damon Runyon Quantitative Biology Fellowship from the Damon Runyon Cancer Research Foundation (DRQ-13-22), the Mahan Fellowship at Herbold Computational Biology Program of Fred Hutchinson Cancer Research Center, the Translational Data Science Integrated Research Center New Collaboration Award and Integrated Research Center at Fred Hutchinson Cancer Research Center, Pilot Award of Immunotherapy Integrated Research Center, and in part through the NIH/NCI Cancer Center Support Grant P30 CA015704. We wish to thank John Heath and Dr. Alphonsus Ng, and all the colleagues from the Institute for Systems Biology for the technical support. Schematic figures in Figure 1, 2d, 3a, 4a, 5a, 6a, 7a were created with BioRender.com, released under BioRender Academic Publication Licenses.

## Author Contributions Statement

Conceptualization, J.X., Y.S., and J.R.H.; Resources, H.C., J.D.G., and J.R.H.; Methodology, J.X., D.G.C., and J.R.H.; Investigation, J.X., D.G.C., W.C., R.N., R.Z., D.Y., J.C., M.K., P.T., B.S., L.J., A.W., Y.R., S.L., R.E., S.H., K.M.M, J.K.L, N.M.F, C.G.L, B.P, H.A., J.W., A.T.M., K.W., P.M., P.D.G., H.C., J.D.G., Y.S., and J.R.H.; Formal analysis, J.X., and J.R.H.; writing – original draft preparation, J.X., and J.R.H.; writing – review & editing, J.X., D.G.C., W.C., R.N., R.Z., D.Y., J.C., M.K., P.T., B.S., L.J., A.W., Y.R., S.L., R.E., S.H., K.M.M, J.K.L, N.M.F, C.G.L, B.P, H.A., J.W., A.M., P.M., P.D.G., H.C., J.D.G., Y.S., and J.R.H.

## Supplementary Figures

Supplementary Fig. 1: Technical validation and methods for antigen and patient assignment

a. Heatmap where each row is a Hashtag used for each SCT-dextramer pool and each column is a single cell. Legend at bottom.

b. Scatter plots of antigen assignment for representative CMV and SARS-CoV-2 antigens. X-axis is the SCT intensity, defined by numbers of UMIs mapped to the SCT-dextramer. Y-axis is the SCT dominance, defined by percent of the cell’s SCT-dextramer associated UMIs that mapped to this antigen’s SCT-dextramer. Cells assigned with the antigen are shown in red positive zone, and have SCT intensity >25, and SCT dominance >25%.

c. Patient assignment by comparison of the WGS SNPs and derived de novo SNPs derived from scRNA-seq data, exampled by Donor-58 and Donor-194. Each panel represents the comparison for each chromosome. Allele frequency and nucleotide identity of reference and alternate are shown in lower right legend.

d. Heatmap with rows as sex specific genes (RPS4Y1 for male, XIST for female), columns are assigned participants. Upper row indicates patient sex from clinical records with blue as male and pink as female. Legend on upper right.

Supplementary Fig. 2: SARS-CoV-2 SCT expression

a. Left: Number of constructed SCT plasmids for putative peptides. Middle: Expressed SCT constructs with useable protein expression yield that were constructed as DNA-barcoded SCT dextramers during 10X Chromium experiment. Note that two of the expressed SCTs were not included due to low sample volume. Right: SCTs that captured SARS-CoV-2 CD8 T cells from COVID-19 participants.

b. Immunodominant A*02:01 epitopes among individuals detected by SCT. Left: CD8+ T cell distribution for each antigen specificity (rows) identified from individual COVID-19 infected participants (each color represents a different participant). Right: total cell counts for each antigen (log10 scale). Only antigens assigned with more than 5 cells were plotted.

Supplementary Fig. 3: TCR Validation via in-vitro Lenti-virus transduction

a. Flow cytometry gating strategy.

b. Selected SCT pMHC tetramer assay on untransduced or TCR-transduced Jurkat cells. TCR#1, previously published by Chour et.al., provides a positive control. We further validate TCR#2 and #3, which are identified from this dataset.

Supplementary Fig. 4: Agreement between SCT expression and NetMHCPan prediction

a. Bar plots comparing expressed vs non-expressed SCT constructs. X-axis: SCT expression status. Y-axis: Normalized SCT expression yield (left) and NetMHCpan predicted binding affinity between peptide and HLA-A*02:01 (right).

b. Scatter plot for normalized SCT expression yield (Y-axis) and predicted binding affinity (X-axis) for all expressed SCTs, each dot represents a SCT construct.

c. Bar plots comparing three groups of peptides based on NetMHCPan prediction output. Y-axis: average hydrophobicity and polarity of peptides within each prediction group.

d. For predicted strong binders, a comparison of the average hydrophobicity and polarity for the ones that showed no/low SCT expression, and the ones showed successful SCT expression.

e. Pie chart showing the SCT expression percentage for peptides in each prediction group. Total peptide number were shown in the top.

For bar plots, data are presented as mean values +/- SEM. The Statistical significance was determined using the two-sided Mann-Whitney U test, and p values are annotated on all relevant plots with exact p-values provided unless p < 0.0001.

Supplementary Fig. 5: Characteristics of Pep-UMAP and Pep-Groups

a. Pep-UMAP colored with selected physicochemical properties, including anchor hydrophobicity, hydrophobicity of the 6^th^ residue, average hydrophobicity, bulkiness, polarity, and absolute charge of exposed (Exp) residues.

b. UMAP embedding of Pep-Groups.

Supplementary Fig. 6: Characteristics of GEX-UMAP

a. UMAP embedding of gene expression (GEX-UMAP) for SARS-CoV-2 specific CD8 T cells color-coded by expression levels of selected mRNA transcripts, and UMAP leiden groups.

b. Dot plot showing normalized expression levels of selected marker genes in each T cell phenotype. The size and color of each dot represent the fraction of expressing cells and the mean of normalized expression levels in each phenotype.

c. Cluster map of phenotype-related genes and surface protein measured via scCITE-seq from Su et. al (2022)^25^. Values are row-normalized, legend on the right. Proteins were marked as red triangle next to surface marker’s name while mRNAs were unlabeled.

d. Stacked bar plot of clone size distribution for T cells captured by threes PG-group with legend on the right.

Supplementary Fig. 7: Characteristics of TCR Groups

a. GEX-UMAP colored with selected physicochemical properties of the TCRβ sequences.

b. Stacked bar plot of clone size distribution for each of the T cell phenotypes with legend on the right.

c. Distribution of CDR3βmer hydrophobicity (top), CDR3β length (middle), CDR3βmer absolute charge (bottom) for each of the TCR-Groups. Bar plots for the respective property are on the right.

d. Bar plots for the mRNA levels of LAG3 and TIGIT.

For bar plots, data are presented as mean values +/- SEM. The Statistical significance was determined using the two-sided Mann-Whitney U test, and p values are annotated on all relevant plots with exact p-values provided unless p < 0.0001.

Supplementary Fig. 8: Validation after removal of dominant TCR clones

a. CD8+ T Cell clonotype distribution for each antigen specificity (rows) (for each antigen, each color represents cells expressing a unique TCR). Only top 15 were plotted.

b. Bar plots evaluating TCR-Groups after removal of large clones. Y-axis: average expression level of representative genes and scores related with Cytotoxicity (top) and Naïve-Memory (bottom). Red (HPhobic-High) and black (HPhobic-Low) bars represent cells within each TCR-Group. Legend on the right. For bar plots, data are presented as mean values +/- SEM. The Statistical significance was determined using the two-sided Mann-Whitney U test, and p values are annotated on all relevant plots with exact p-values provided unless p < 0.0001.

Supplementary Fig. 9: Characteristics of PG-TCR groups

a. Radar graphs of physicochemical properties for example PG-TCR groups. Blue and grey shaded areas of the outer rings indicate Pep-Group, and TCRβ properties, respectively. Each axis displays the normalized average value for each property, with lowest value in the center. The shaded polygons reflect the property space occupied by the peptide-TCRβ groupings. Legend on the bottom.

b. GEX-UMAP color encoded with densities of PG-TCR groups based on each cell’s antigen specificity and TCR sequence.

c. Stacked bar plot of phenotype distribution for each of the PG-TCR groups with legend on the right.

d. Surface protein validation with respect to findings in Main Fig. 6. Heatmap with X-axis as PG-TCR group assignment and Y-axis as level of a given protein normalized per row and column, see legend on right.

e. Stacked bar plot of PG-TCR group distribution for acute and post-acute timepoint with legend on the right.

f. Clustermap of cells by expression levels of selected mRNA transcripts for each timepoint. Legend on the right.

g. Bar plots comparing expression levels of selected mRNA transcripts for cells at acute and post-acute timepoint. Number of cells for each group: PG3:High Acute (n=448), Post-Acute (n=35); PG1: Acute (n=116), Post-Acute (n=156). For bar plots, data are presented as mean values +/- SEM. The Statistical significance was determined using the two-sided Mann-Whitney U test, and p values are annotated on all relevant plots with exact p-values provided unless p < 0.0001.

Supplementary Fig. 10: TCR alpha chain analysis

a. Analysis of which physicochemical characteristics of the TCR alpha chain exhibit significant associations with T cell phenotype. The top differential TCR CDR3α physicochemical properties for effector (Cytotoxic, EM and Hybrid) phenotypes, relative to non-effector (CM and naïve) phenotypes. The Statistical significance was determined using the two-sided Mann-Whitney U test, and p values are annotated on all relevant plots with exact p-values provided unless p < 0.0001.

Supplementary Fig. 11: External dataset validation

a. Bar plots comparing expression levels of selected mRNA for cells within each TCR- Groups. Number of cells in each group: HPhobic-High (n = 1841), HPhobic-Low (n = 1988). Original dataset from Fischer et. al (2021)^58^. For bar plots, data are presented as mean values +/- SEM. The Statistical significance was determined using the two-sided Mann-Whitney U test, and p values are annotated on all relevant plots with exact p-values provided unless p < 0.0001.

b. Pearson correlation coefficients between selected mRNA levels and TCR CDR3β properties for CMV-NLVP (left) and Influenza-GILG (right) specific CD8 T cells. Original dataset from Chen et. al (2023)^8^.

## Supplementary Tables

Table S1: Clinical characteristics, medical history of INCOV sub cohort in this study Table S2.1: List of putative SARS-CoV-2 peptide antigens for SCT-pMHC expression Table S2.2: Vireo output for SNP demultiplexing

Table S2.3: TCR gene usage for top 5 SARS-CoC-2 epitopes

Table S2.4: List of TCRs used for validation (via lenti-virus transduction and SCT-tetramer binding assay)

Table S3: Amino acid property scales used for peptide and TCR residues Table S4: Pep-UMAP characteristics

Table S5: Filtered DEGs in cells assigned to PG3 compared to PG2 antigen

Table S6: Enriched pathways (Reactome 2022) for upregulated genes in PG3 vs PG2 Table S7: Log2 fold change of TCR physicochemical properties, effector phenotypes vs non-effector phenotypes

Table S8: Mann-Whitney U test of TCR physicochemical properties, effector phenotypes vs non-effector phenotypes

Table S9: DEGs in cells in HPhobic-High compared to HPhobic-Low

Table S10: Enriched pathways (Reactome 2022 and GO Biological Process 2023) for upregulated genes in HPhobic-High vs HPhobic-Low

Table S11: Cell count at Acute and Post-Acute time points

