## Supplementary figures for "APMAT analysis reveals the association between CD8 T cell receptors, cognate antigen, and T cell phenotype and persistence"

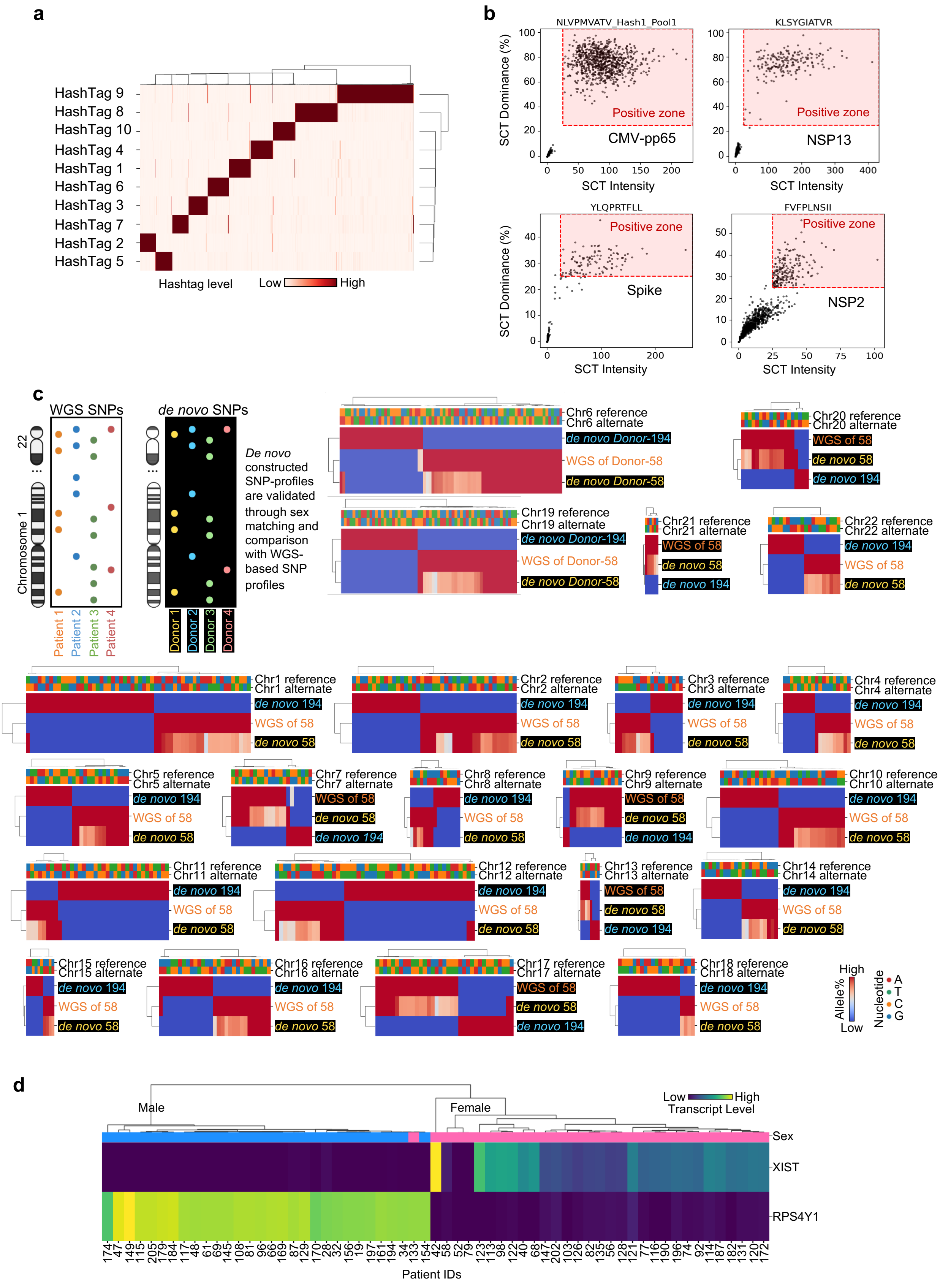


Supplementary Fig.1

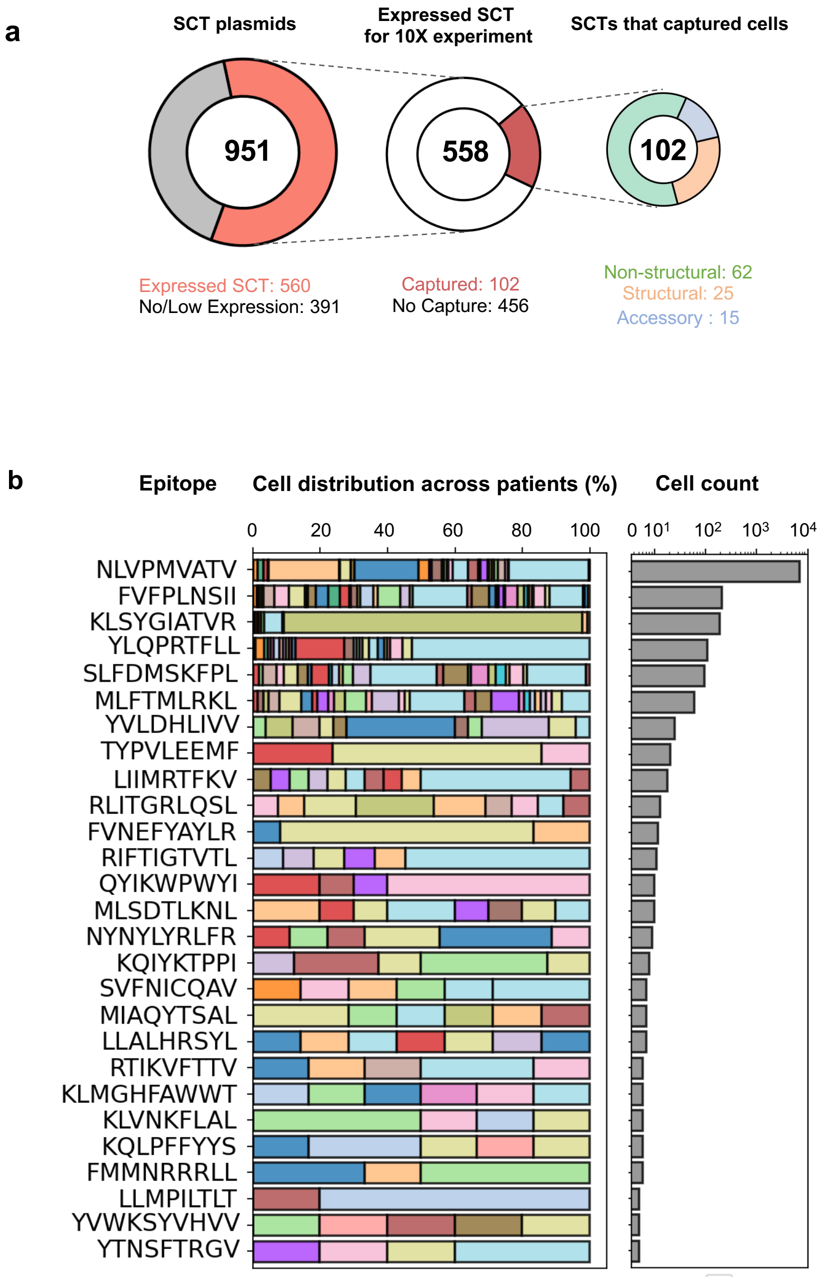


Supplementary Fig.2

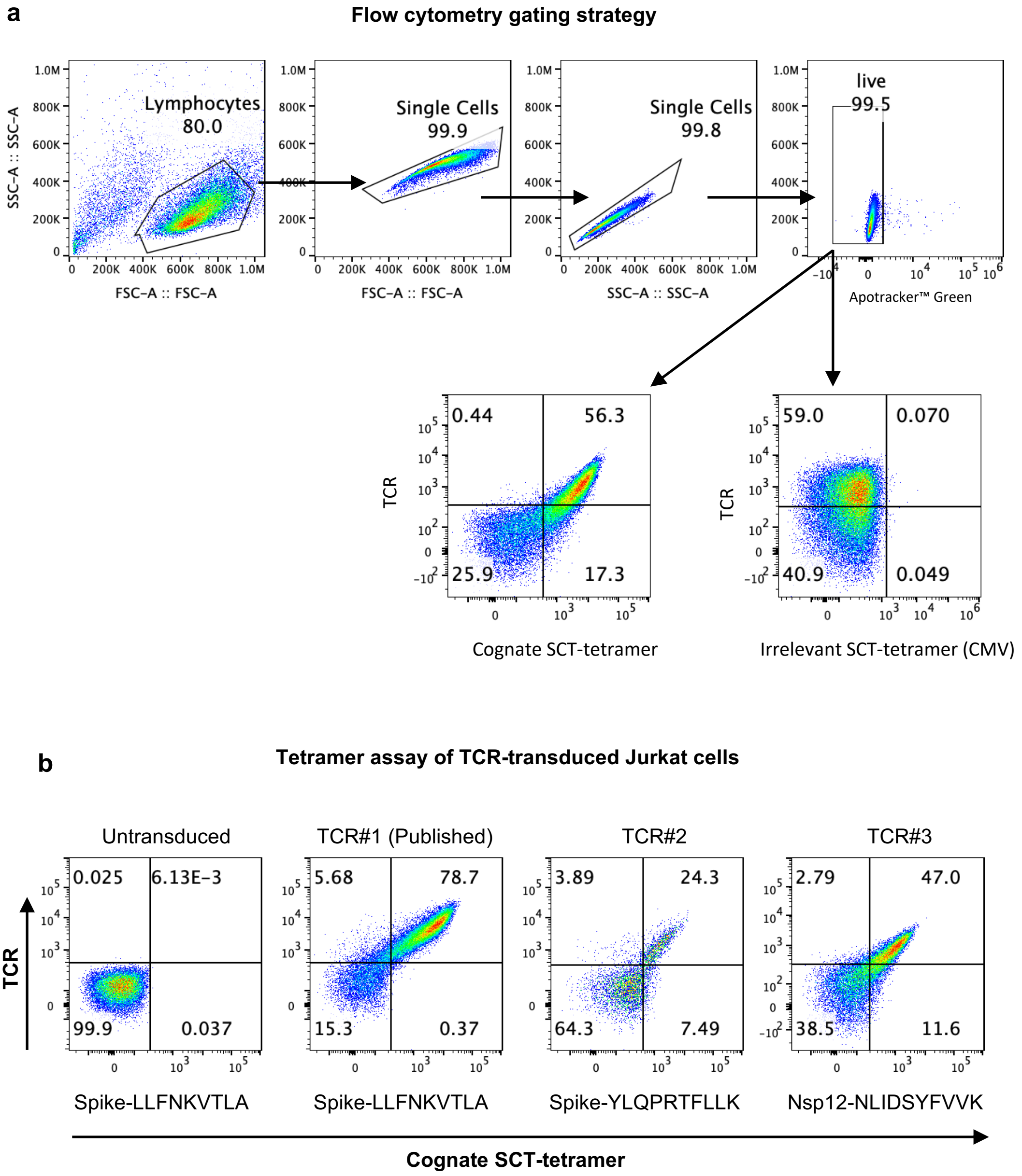


Supplementary Fig.3

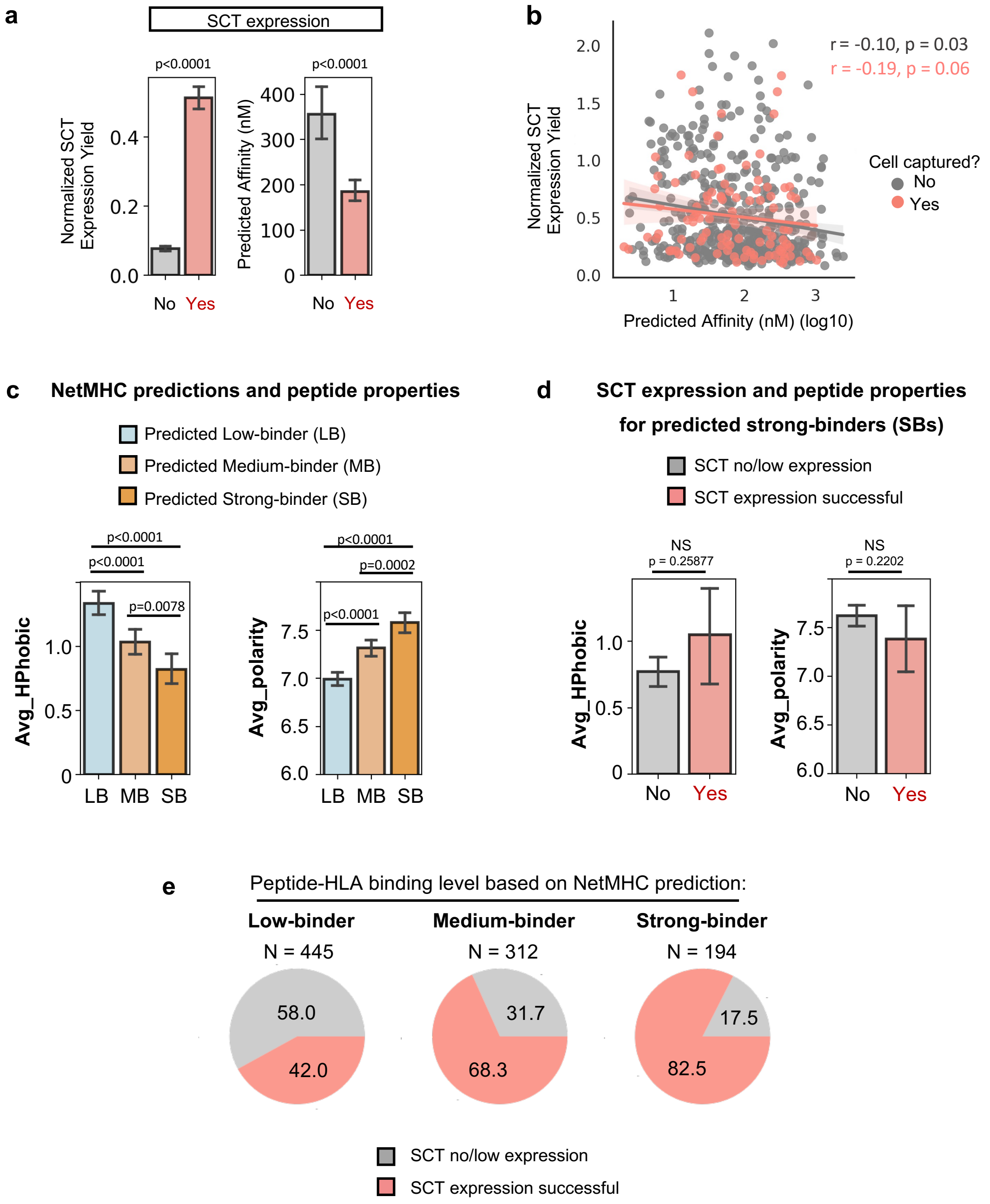


Supplementary Fig.4

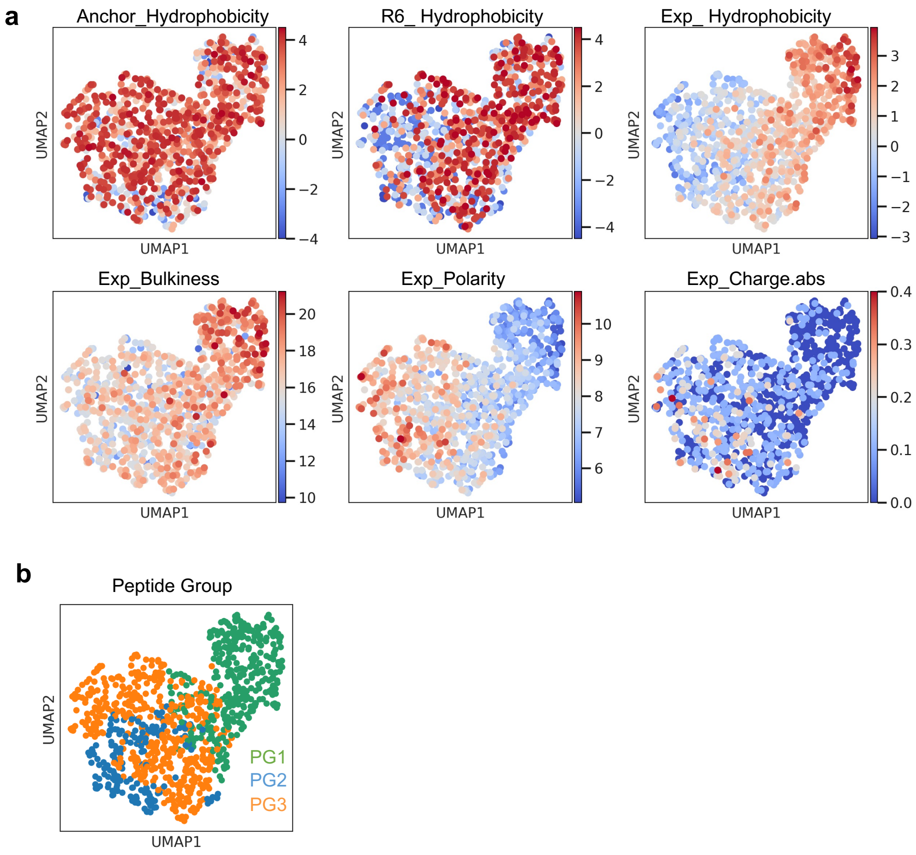


Supplementary Fig.5

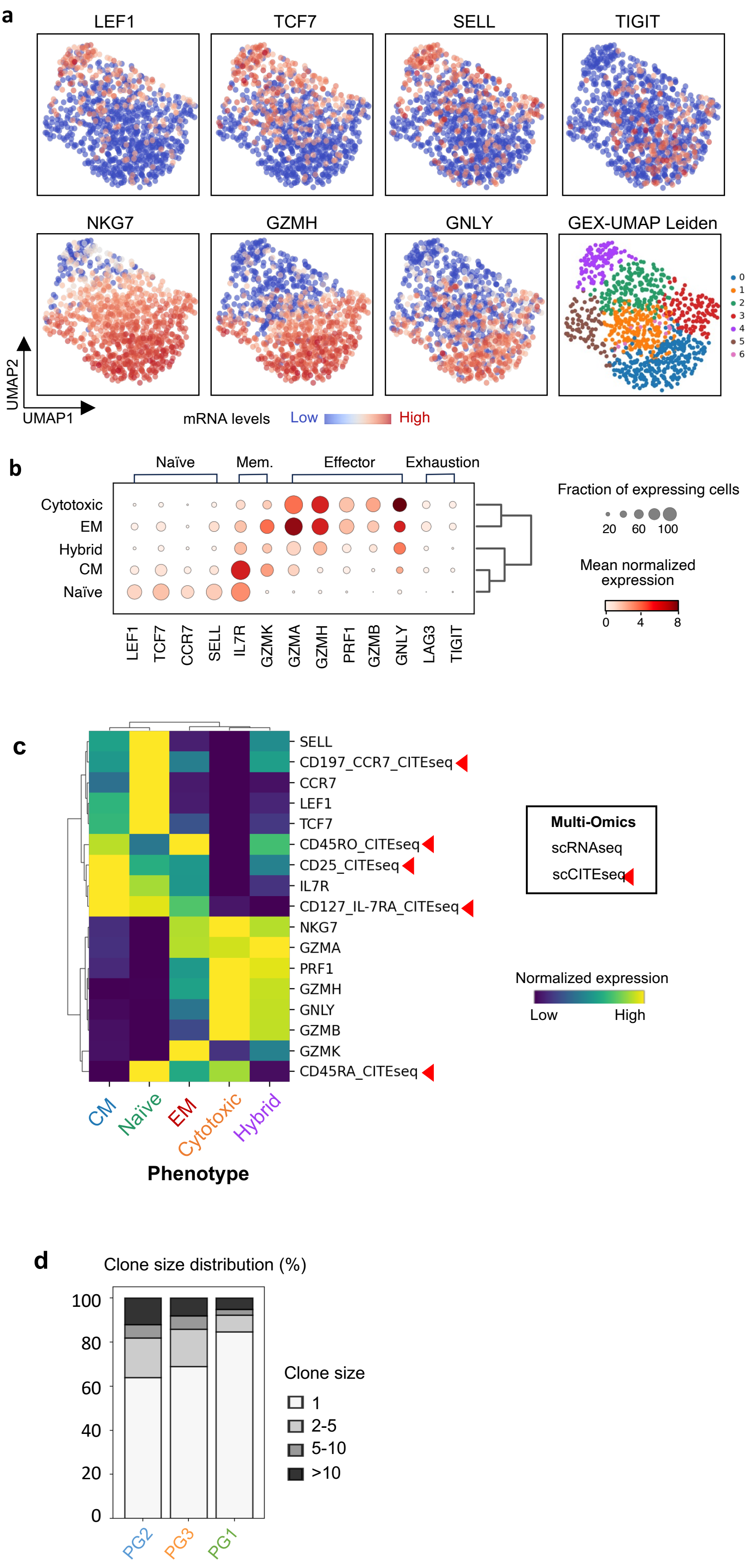


Supplementary Fig.6

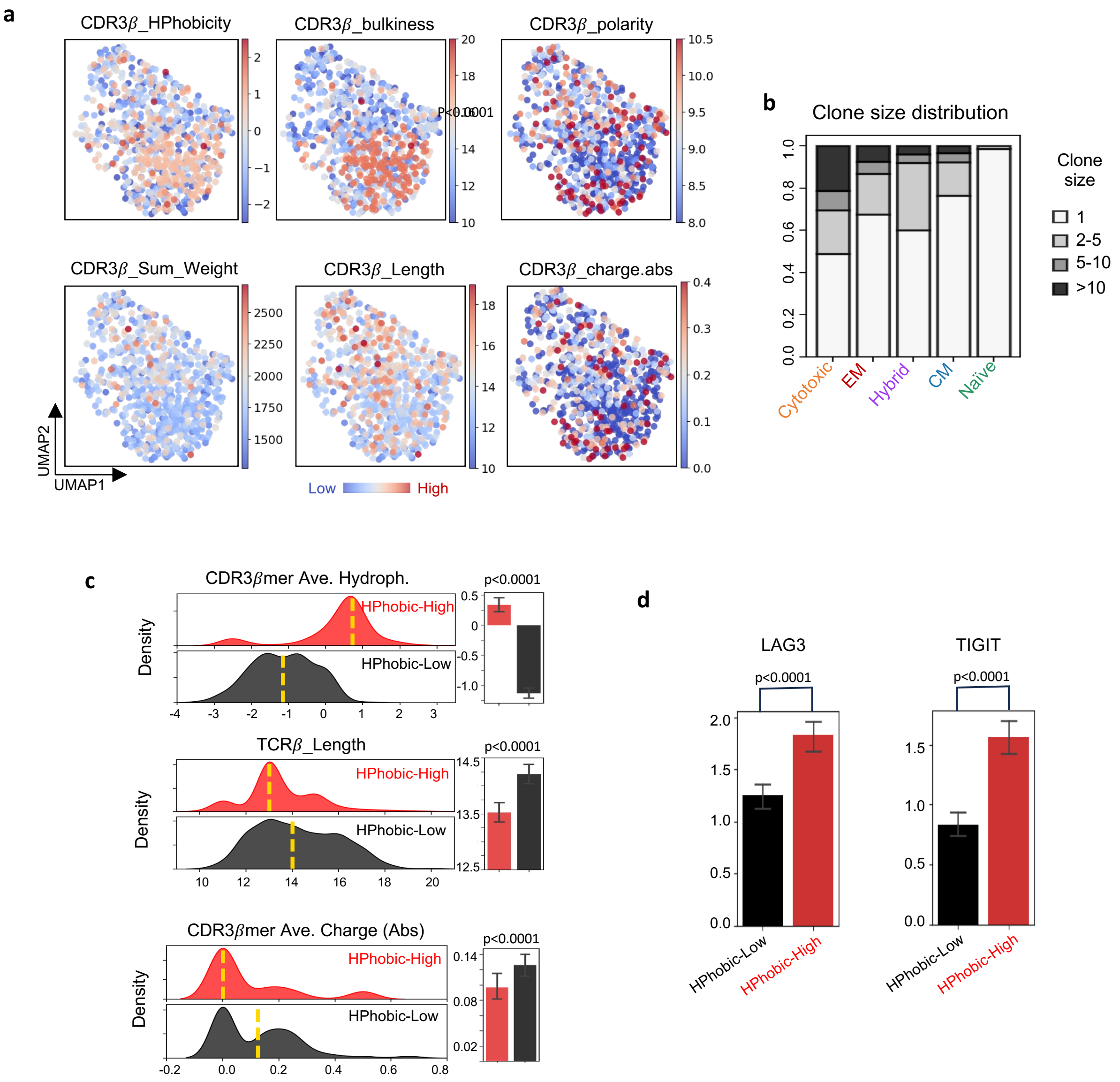


Supplementary Fig.7

**a**

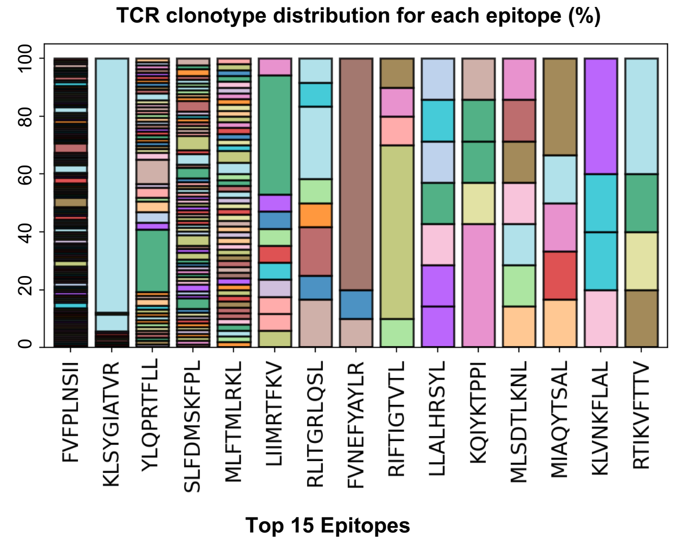


**b**

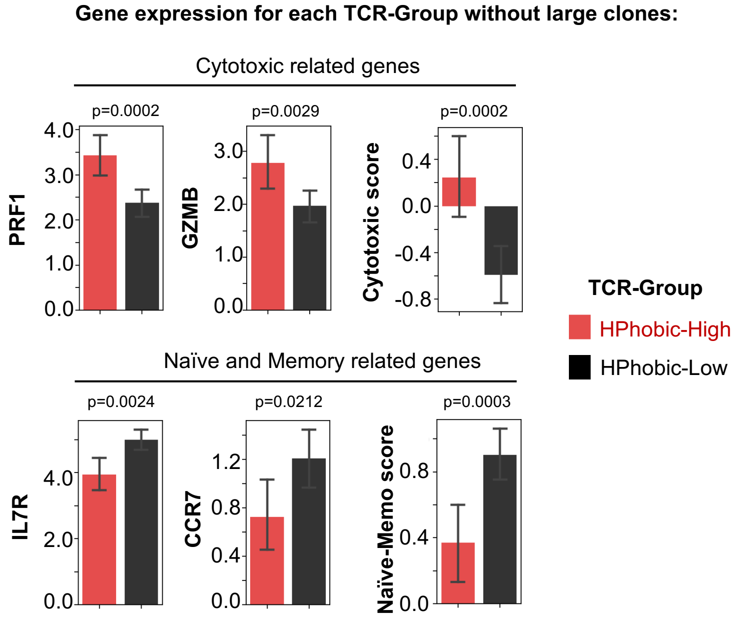


Supplementary Fig.8

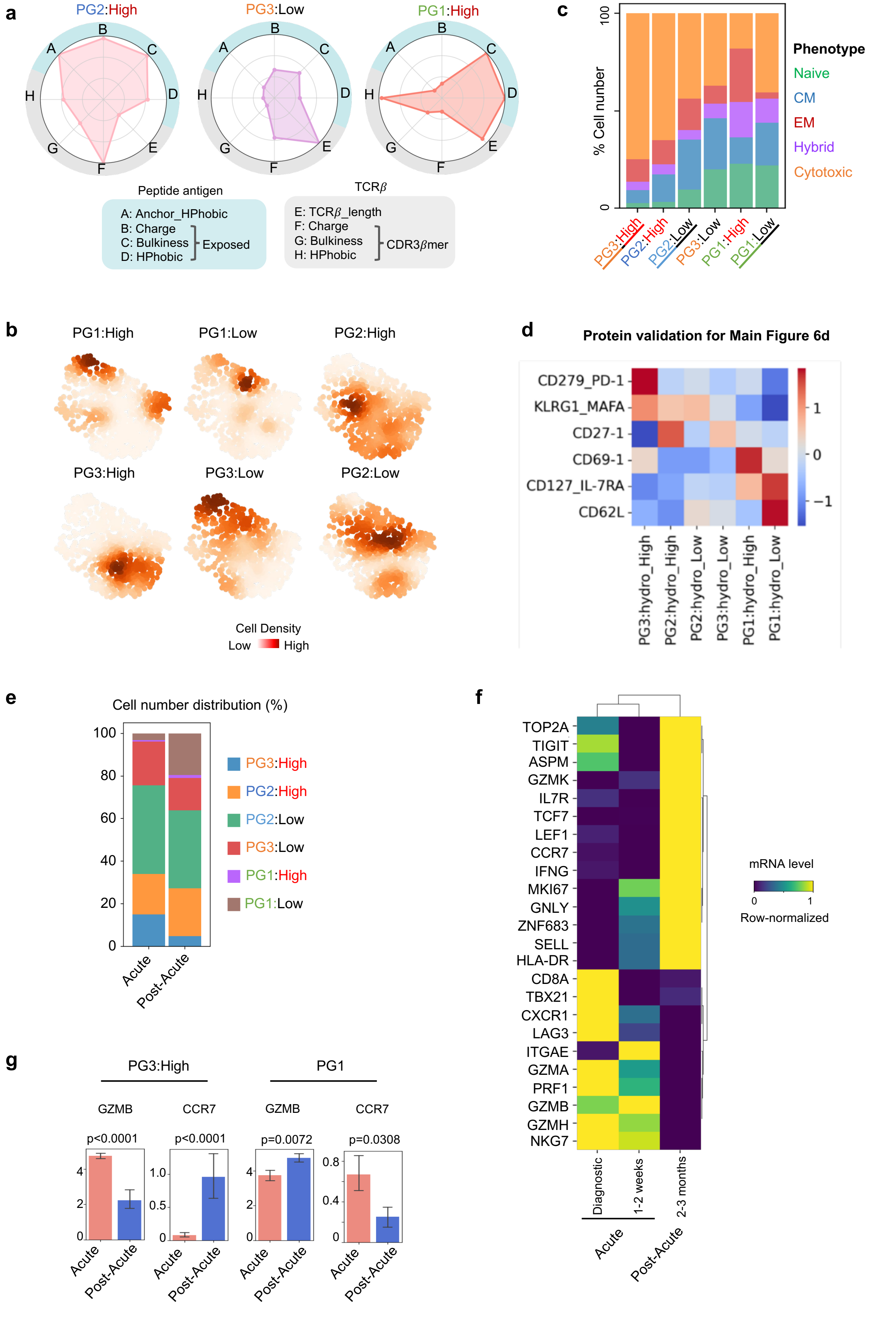


Supplementary Fig.9

**a**

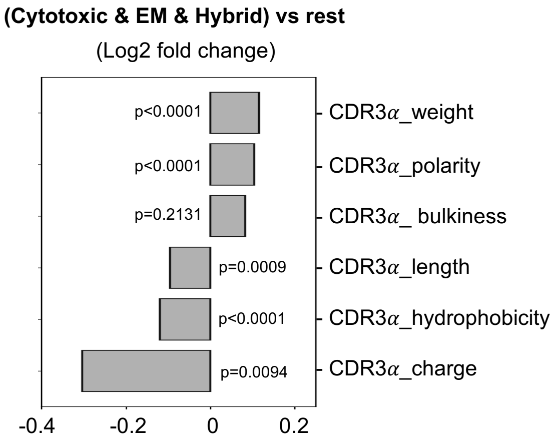


Supplementary Fig.10

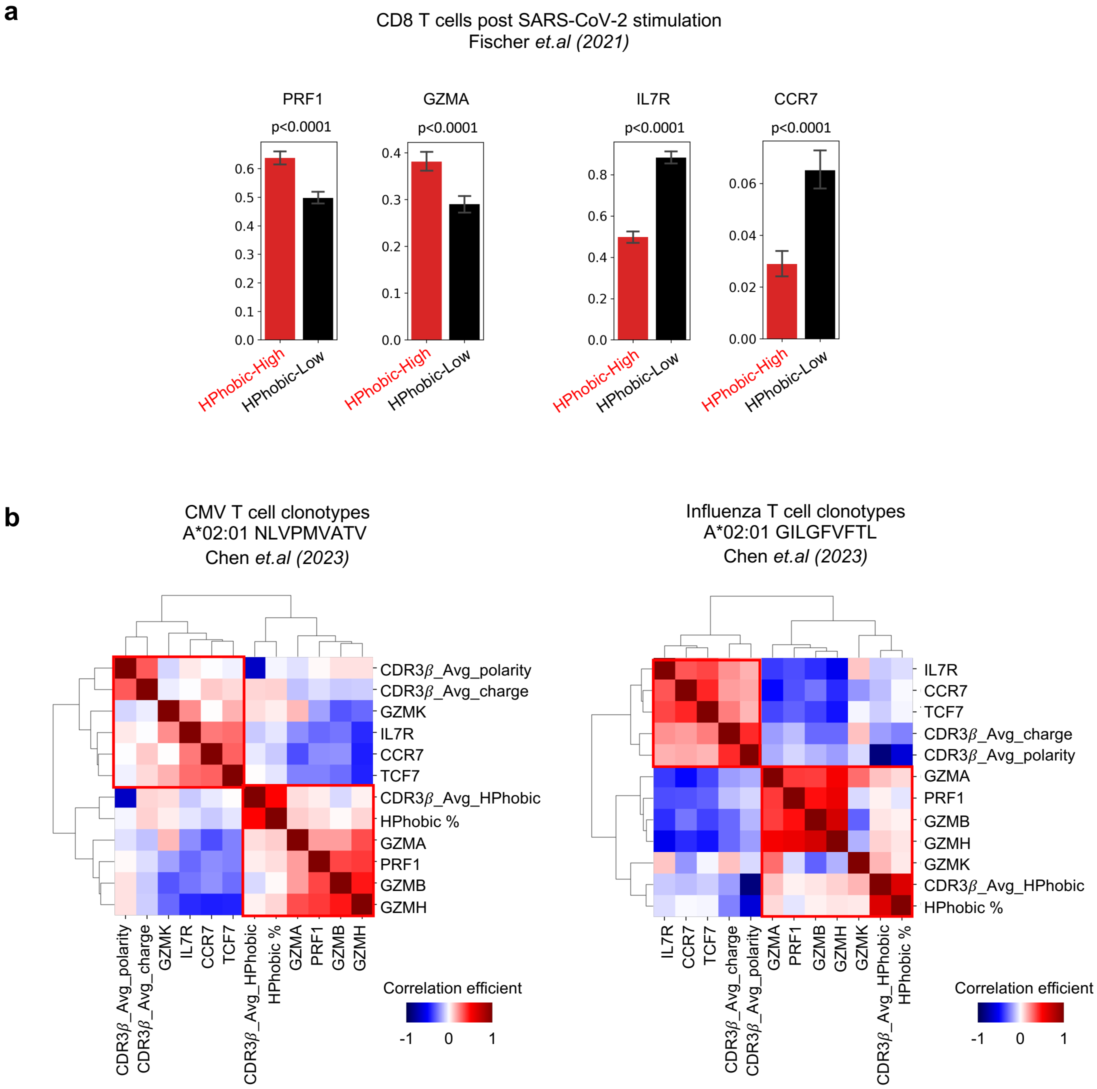


Supplementary Fig.11
